# The Impact of Uremia and Intestinal Dysbiosis on Hepatic Drug Metabolism in a Rat Model of Progressive Chronic Kidney Disease

**DOI:** 10.1101/531939

**Authors:** Emily D Hartjes, Yong Jin Lim, Thomas J Velenosi, Kait F Al, Jean M Macklaim, Andrew S Kucey, Gregor Reid, Jeremy P Burton, Gregory B Gloor, Bradley L Urquhart

**Affiliations:** Department of Physiology and Pharmacology, Schulich School of Medicine and Dentistry, Western University, London, Ontario, Canada; Department of Biochemistry, University of Western Ontario, London, Ontario, Canada N6A 5B7; Department of Microbiology & Immunology, Western University, London, Canada; Canadian Centre for Human Microbiome and Probiotics, London, Canada; Department of Medicine, Schulich School of Medicine and Dentistry, Western University, London, Ontario, Canada

**Keywords:** Uremia, gut microbiota, dysbiosis, cytochrome P450, drug metabolism, disease progression, metabolomics, sequencing

## Abstract

Nonrenal clearance pathways such as drug metabolism are decreased in chronic kidney disease (CKD). Although the mechanism remains elusive, uremic toxin retention and an altered gut microbiota are suspected to influence cytochrome P450s (CYPs) contributing to the unpredictable pharmacokinetics in patients with CKD. We characterized dysbiosis and uremia in CKD to elucidate associations between CYP expression and CKD progression. Rats fed control or CKD-inducing adenine diet were subsequently studied at five time points over 42 days. CYP expression and activity were compared to alterations in the 1) plasma and liver metabolome and 2) gut bacterial microbiota. CYP3A2 and CYP2C11 were downregulated in CKD by ≥76% (p<0.001) concurrently with or slightly prior to CKD onset as defined by serum creatinine. Metabolite profiles were altered prior to changes in the gut microbiota, and gut-derived uremic toxins including indoxyl sulfate, phenyl sulfate and 4-ethylphenyl sulfate correlated with CYP3A2 or CYP2C11 expression. Bacterial genera *Turicibacter* and *Parabacteroides* were identified as being characteristic of CKD. In conclusion, CYP3A2 and CYP2C11 are downregulated before dysbiosis and correlate with select uremic toxins.

## Introduction

Chronic kidney disease (CKD) is a progressive and irreversible loss of kidney function over time affecting approximately 16% of the global population (1). Progressive loss of kidney function in CKD leads to accumulation of waste products in plasma, resulting in uremia (2). Uremia contributes to the progression of CKD into end-stage renal disease (2) and has also been associated with the activation of the immune response, gut microbial alterations (3), and cardiovascular events (4).

The impairment of renal drug clearance in CKD is well established. However, non-renal, hepatic drug clearance is also significantly decreased in CKD. Non-renal drug clearance was identified by KDIGO (Kidney Disease: Improving Global Outcomes) as an important consideration for dose recommendations (5). CKD patients experience altered pharmacokinetics, partially mediated by altered activity of cytochrome P450 (CYP) drug metabolizing enzymes (6, 7). CYP3A4 and CYP2C9 metabolize approximately 43% of all clinically relevant drugs (8). Although several studies suggest hepatic drug metabolism is decreased in CKD (6, 7), the exact mechanisms underlying CYP downregulation are poorly understood. Uremia, hormones, gut bacterial alterations and associated inflammation are all factors proposed to affect drug metabolizing enzymes in CKD. Despite CKD being a progressive disease, many studies focus exclusively on severe CKD while the majority of patients suffer from earlier stages of the disease. This leaves a gap in our understanding of the temporal relationship between CKD progression and changes in CYP expression. Uremia may influence CYP enzyme changes by altering transcriptional regulation of CYP enzymes (9); modulating inflammation or parathyroid hormone (PTH) (10) or direct inhibition by uremic toxins (11–13).

The relationship between gut bacteria and host physiology/pathophysiology has been extensively studied. Alterations in gut bacteria have been linked with inflammatory bowel disease, obesity, cardiovascular disease, asthma and cancer (14, 15). Dysbiosis refers to changes in bacterial composition associated with a non-infectious disease state (15). In a large cohort study of 1106 human stool samples, glomerular filtration rate (GFR) was a major factor associated with altered bacterial composition (16). Thus, it comes as no surprise that patients with CKD also exhibit dysbiosis (16).

In this study, we hypothesized that CKD would cause uremia and gut microbial changes detectable prior to downregulation of drug metabolizing enzymes. Bacterial alterations and metabolic changes have not been comprehensively studied with respect to altered drug metabolism throughout the temporal progression of CKD. To our knowledge, this is the first study to use metabolomics and 16S bacterial rRNA gene sequencing to investigate CYP expression and activity over CKD progression. Our study aims to characterize plasma and liver uremic toxins, the gut microbial composition and CYP expression and activity over the progression of CKD.

## Materials & Methods

### Animal Model & Study Design

Sixty-six male Wistar rats (150g) were obtained from Charles River Laboratories, Inc. (Wilmington, MA) and randomized into six groups defined by time of euthanasia (day 0, 3, 7, 14, 28 and 42). Each time point consisted of six control and six CKD rats. Rats were housed with a same-group cage mate to minimize coprophagy alterations of the gut microbiota. Rats were given either 0.5% adenine supplemented chow to induce CKD or standard chow pair fed to match caloric intake. Rats were euthanized by isoflurane anesthetization followed by decapitation. Blood was collected in heparinized tubes and liver was snap-frozen in liquid nitrogen. Caecal samples were obtained on a sterile, single culture swab (BD, Sparks, MD) touched to an open incision of the caecum. All samples were stored at −80°C until further analysis excluding the right kidneys which were stored in 10% formalin.

### Disease Markers & Histology

Conventional CKD markers urea and creatinine were measured in rat plasma using standard methods by the Pathology and Laboratory Medicine group (PaLM, London, ON; www.lhsc.on.ca/palm/). Kidney tissue and histological images were prepared as previously described (17). Light microscopy and photographs of prepared haematoxylin and eosin stained slides were obtained on a Leica DM1000 light microscope paired with a Leica DFC295 camera and Leica Application Suite v3.8.0 software.

### Real-Time Polymerase Chain Reaction (PCR)

Total mRNA was extracted from rat liver, tested for purity and quantified using quantitative PCR with methods and validated primers as stated previously (9). Gene expression was normalized to β-actin using the ΔΔCT method.

### Western Blotting

Hepatic microsomal fractions were prepared by differential centrifugation as previously described (17). Western blot analysis was performed as previously published with minor alterations (17) and all blots were completed in duplicate for both CYP3A2 and CYP2C11.

### Enzymatic Activity

Enzymatic activity was analyzed by incubating microsomal fractions with testosterone, a known substrate of CYP3A2 and CYP2C11. Testosterone metabolites 6βOH-testosterone (CYP3A2) and 16αOH-testosterone (CYP2C11) were measured via mass spectrometry (MS) in a 96-well plate assay adapted from (17). In a final volume of 75µL, 0.2mg/mL microsomal protein and reaction buffer (50mM potassium phosphate with 2mM MgCl_2_ pH 7.4) was incubated with 1µL testosterone (Steraloids Inc., Newport, RI) at concentrations of 12.5, 25, 75, 200 and 400µM for 10 min at 37°C. All reactions were initiated with the addition of 1mM NADPH (Sigma Aldrich) and shaken at 37°C for 20 min before the reaction was terminated using 225µL ice-cold acetonitrile with 80ng/mL flurazepam internal standard (Cerilliant, Round Rock, TX). Plates were shaken, centrifuged at 4000×g for 10 min, supernatant diluted 5-fold with milliQ water. Enzymatic products were separated on a Phenomenex Kinetex phenyl-hexyl column (1.7µm particle size, 50mm × 2.1 mm) maintained at 40°C in a Waters ACQUITY UPLC I-Class System (Milford, MA). Mobile phase flow was set to 0.5 ml/min and consisted of UPLC-grade water (A) and acetonitrile (B) both containing 0.1% formic acid with a gradient as follows: 0–0.5 mins, 25% B; 0.5–2 mins 25–35% B; 2–2.5 mins 35–80% B; 2.5–3.5 mins held at 80% B; 3.5 mins 25% B. Analytes were detected using quadrupole time-of-flight mass spectrometry (QTof/MS) on a Waters Xevo^TM^ G2S-QTof/MS and Waters ACQUITY I-Class UPLC with parameters as previously described (17). Mass-to-charge ratios for hydroxy testosterone were targeted (*m/z* = 305.2117) for quantification using QuanLynx v4.1 software. Michaelis-Menten curves were generated with GraphPad Prism (v6.0; GraphPad Software Inc., San Diego, CA).

### Untargeted Metabolomics

#### Sample & Batch Preparation

Plasma and liver samples were prepared as previously described (18) with 3:1 ice-cold acetonitrile and 2.5µM chlorpropamide internal standard (Sigma Aldrich) then run on both the Waters ACQUITY UPLC HSS T3 (1.8µm particle size, 100 mm × 2.1 mm) reverse-phase liquid chromatography (RPLC) column and the Waters ACQUITY BEH Amide (1.7µm particle size, 100 mm × 2.1 mm) hydrophilic interaction liquid chromatography (HILIC) column. Supernatant was either diluted 5-fold in water for RPLC or directly injected for HILIC. Sample injection order was randomized, and a quality control sample made from pooled samples was run every ten injections. All samples were run in a single batch for each biological matrix and column.

#### Chromatography & Mass Spectrometry

Columns were maintained at 45°C and mobile phase flow set to 0.45 ml/min consisting of UPLC-grade water (A) and acetonitrile (B), both containing 0.1% formic acid. The RPLC anlysis was run as previously described (18). The HILIC column followed a gradient of 0–0.5 mins 99% B; 0.5–6 mins 99–50% B; 6–8 mins 50–30% B; 8–8.5 mins 30–99% B. Samples were run separately in succession for both positive and negative electrospray ionization (ESI) modes on the UPLC-QTof/MS instrument. Mass spectrometer source, method, calibration and other parameters were identical to those in (18) and data was collected by MassLynx v4.1 software (Waters).

#### Data Processing

Data processing for each run and ionization mode was performed separately in R studio (v3.2.3). MassLynx data files were converted to mzData files using convert.waters.raw package v1.0 (github.com/stanstrup/convert.waters.raw). Pooled samples were used to find the optimal peak picking parameters, retention time corrections and grouping parameters with the isotopologue parameter optimization package v1.0.0 (github.com/rietho/IPO/blob/master/vignettes/IPO.Rmd). The resulting parameters were inputted into the XCMS package v1.50.1 to pick appropriate peaks, integrate the area under the curve and replace zero values (19). The CAMERA package v1.32.0 was used to annotate possible isotopes and adducts (20). XCMS and CAMERA packages were used to integrate positive and negative ionization modes before normalizing to internal standard and applying a threshold of 30% variability of the quality control.

#### Metabolite Identification

The accurate monoisotopic mass (*m/z*) and fragmentation spectrum of each metabolite was used to identify metabolites via METLIN, MassBank or Human Metabolome Database (HMDB) (21). Metabolites were identified in accordance with the reporting standards for metabolite identification (22).

### In Vitro Assessment of Uremic Toxins on CYP3A4 Expression

Human hepatoma Huh7 cells were maintained in Dulbecco’s modified Eagle’s medium supplemented with 10% FBS, 100 IU/ml penicillin, and 100 µg/ml streptomycin and 2 mM L-glutamine. Cells were grown at confluence for 4 weeks prior to treatment to ensure adequate CYP3A4 expression levels (23). Huh7 cells were treated with select uremic toxins: creatinine (2121.6 µM), *p*-cresyl sulfate (186.1 µM), indoxyl sulfate (1113.2 µM), urea (76.6 mM) for 24 hours. An indoxyl sulfate concentration-response effect on CYP3A4 mRNA expression was generated using a concentration range of indoxyl sulfate found in normal and CKD patients (0 to 1000 µM) with 40 g/L human serum albumin (HSA, Lee Biosolutions, St. Louis, MO). Indoxyl sulfate was incubated in media containing 40 g/L HSA for 3 hours at 37°C to allow for plasma protein binding equilibration prior to cell treatment. Cell viability was assessed using the TACS 3-(4,5-dimethylthiazol-2-yl)-2,5-diphenyltetrazolium bromide (MTT) assay as previously described (24).

### Gut Microbial Sequencing

#### Illumina Sequencing

DNA was extracted from caecum swabs using the PowerSoil-96 Well DNA isolation kit from MoBio using convenience modifications of the Earth Microbiome Project protocol (25). 288 unique primer combinations were established using the 515F and 806R barcoded primers (25, 26) to amplify the V4 variable region of the 16S rRNA gene. Primers followed the template: Forward primer [5’-ACACTCTTT CCCTACACGACGCTCTTCCGATCTnnnn(8)GTGCCAGCMGCCGCGGTAA-3’] and reverse primer [5’-CGGTCTCGGCATTCCTGCTGAACGCTCTTCCGATCTnnnn (8)GGACTACHVGGGTWTCTAAT-3’] where the 5’ end is the Illumina adaptor sequence, the nnnn indicates four random nucleotides, (8) represents one of 36 barcoded sequences and the 3’ end is the primer region for V4 (Supplementary Table 1). Amplification was carried out in 42µL total volume with 20µL primer mix (3.2pmol/µL per primer), 20µL GoTaq Hot Start Mastermix (Thermo Scientific) and 2µL template DNA then run for 2 min at 95°C followed by 25 cycles of 1 min at 95°C; 1 min at 52°C and 1 min at 72°C excluding a final elongation. Barcoded PCR products were quantified with a Qubit dsDNA assay kit on a Qubit 2.0 (Life Technologies), normalized by amount of DNA, pooled then purified with a PCR clean-up column. The cleaned DNA was amplified once more with primers OLJ139 [5’AATGATACGGCGACCACCGAGATCTACACTCTTTCCCTACACGA3’] and OLJ140 [5’CAAGCAGAAGACGGCATACGAGATCGGTCTCGGCATTCCTGCTG AAC3’] before paired-end sequencing on the Illumina MiSeq platform at the London Regional Genomics Centre (LRGC, lrgc.ca, London, ON).

#### Data Processing

Paired reads, each 220bp long were processed with the Illumina_SOP protocol accessed through Github with minor convenience revisions (https://github.com/ggloor/miseq_bin). After demultiplexing, raw reads were overlapped with a minimum 30 nucleotides using Pandaseq (v2.5) (27) then filtered with in-house Perl and UNIX scripts to ensure exact barcode matching and primer matching with up to two allowable mismatches (28). OTUs were clustered at 97% identity using the uSearch (v7.0.1090) tool (29) and the most abundant sequence in the OUT was annotated via the mothur script (30) to search the Silva 16S rRNA gene reference database (Silva.nr_v119) (31, 32). In mothur, a bootstrap cut-off of 70% was used for taxonomical identification and redundancy. A total of 1199 OTUs were retained across all samples (Supplementary Table 2). In R studio (v3.2.3) the zCompositions (v1.0.3-1) package (33) was used for zero-replacement before data was centered-log ratio (clr) transformed for compatibility with downstream multivariate statistical analysis (34, 35).

### Statistical Analysis

#### Disease Markers, Real-Time PCR, Western Blotting & Enzymatic Activity Assay

CYP measurements and disease markers urea and creatinine are presented as mean ± SEM and analyzed by 2-way ANOVA paired with Sidak’s multiple comparisons test. \**p* < 0.05 compared to matching day control indicates significance.

#### Untargeted Metabolomics

MassLynx software and the EZInfo v2.0 package (Umetrics, Umeå, Sweden) were used to perform principal component analysis (PCA) to evaluate the initial separation between CKD and control over time for each of the four analytical runs. Data was Pareto scaled and pooled samples were run to confirm minimal variance. Multivariate analysis was performed on each day of each run. EZInfo was used to generate orthogonal partial least squares discriminant analysis (OPLS-DA) of the original PCA (Supplementary Figure 1). To assess multivariate OPLS-DA sufficiency, each comparison received a goodness of fit value ratio threshold (R^2^/Q^2^ < 2) (36). Subsequent thresholds were applied (VIP > 0.8; p(corr)[1] > 0.4 or < −0.4) by finding the variable importance in projection (VIP) and the p(corr)[1] axis as a measure of magnitude and difference between treatments (37) (Supplementary Figure 1). Only metabolites that met or exceeded the thresholds on two or more consecutive time points were retained for comparison with univariate and correlative analyses. Univariate analysis was performed by the open-access online software MetaboAnalyst 3.0 to conduct a p-value corrected (FDR = 0.05) independent 2-way ANOVA on each metabolite via the “Time series” and “Two-factor independent samples” applications (21). Significance (p<0.05) was required for both “*Time*” and “*Disease*” to retain the metabolite for comparison with multivariate and correlative analyses. Spearman correlations were conducted between each metabolite and the mRNA, protein or enzymatic activity levels of each enzyme. Metabolomics datasets were matched by sample to the corresponding CYP dataset and correlation coefficients (r-values) manually filtered with high stringency (r > 0.65 or r < −0.65). Metabolites that did not also satisfy univariate analysis were removed from the correlation subset. Metabolites that did not satisfy multivariate analysis are indicated but retained to capture biologically relevant changes independent of magnitude.

#### Caecal Microbiota

Multivariate PCA was performed in EZInfo as described above for untargeted metabolomics excluding scaling. To evaluate univariate differences between CKD and control groups, the effect size and overlap for each bacterial taxonomic group was calculated for each time point individually using the R package ALDEx2 (v1.2.0) (bioconductor.org/packages/release/bioc/html/ALDEx2.html) (38, 39). Severe thresholds were applied to both effect size (> 1.5 or < −1.5) and overlap (< 6.5%) for each bacterial abundance (39, 40). Significance was defined as satisfying the effect size and overlap thresholds with 95% confidence. Species and strain information were manually searched using the Targeted Loci Nucleotide BLAST application through NCBI (blast.ncbi.nlm.nih.gov/Blast.cgi).

## Results

### Model of CKD Progression

CKD markers urea and creatinine both showed significant increase in CKD rat plasma by day 14 (Figure 1 A-B). This increase continued to a 9-fold and 11-fold difference between CKD and control for urea and creatinine, respectively, on day 42. Kidney histology showed enlarged tubules, inflammation and fibrosis by day 14 through to day 42 (Figure 1 C-H). Animal weights did not change between groups (Supplementary Table 3).

**Figure 1.**
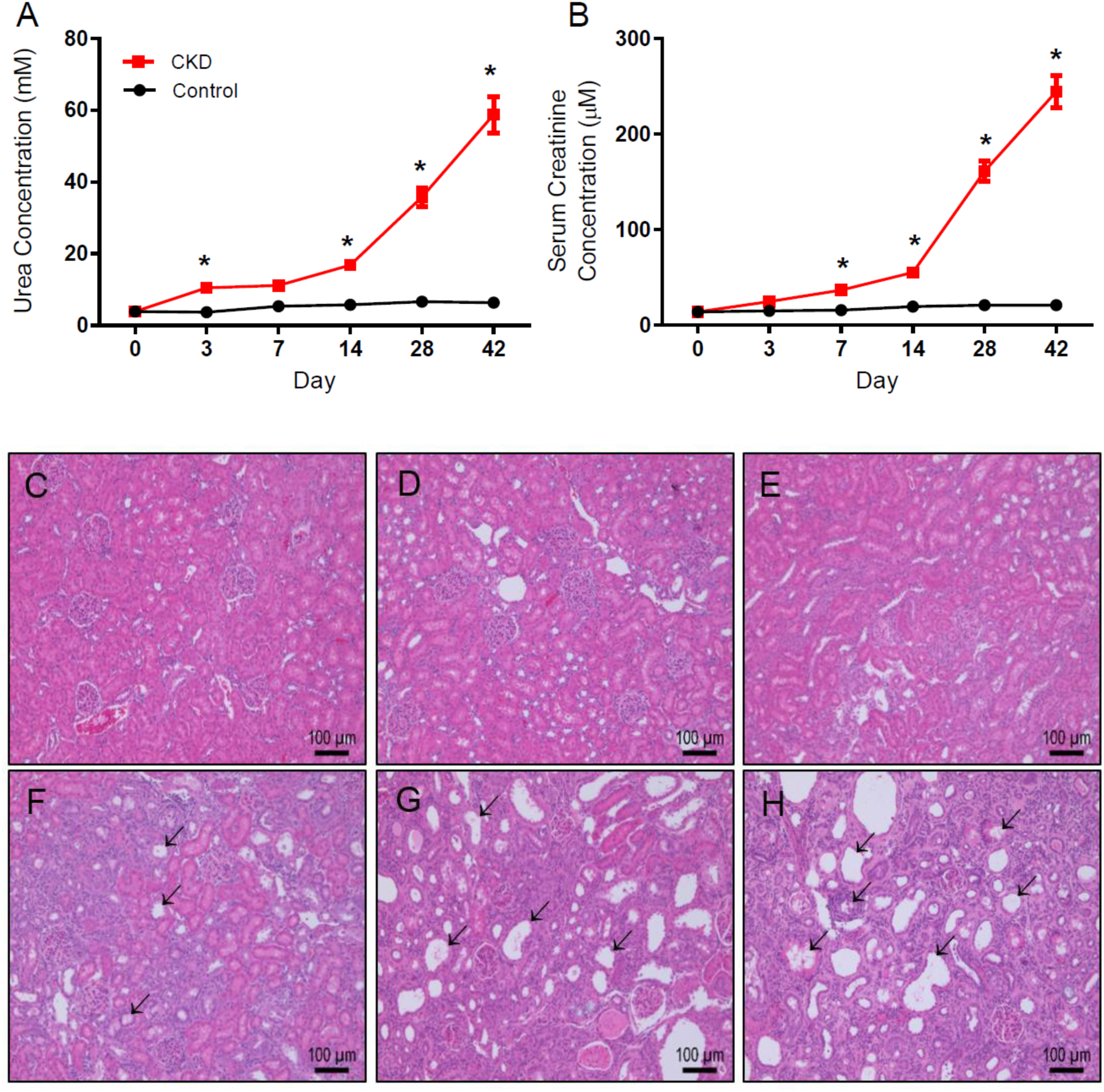
Assessment of CKD in Wistar rats orally administered 0.5% adenine over 42 days. (A) Plasma urea (mM) and (B) serum creatinine (µM) concentrations of control and CKD rats presented as mean ± SEM. \**p* < 0.05 when compared to matching day control; n ≥ 6. H&E stained rat kidney sections from day 0 control (C) and CKD days 3 (D), 7 (E), 14 (F), 28 (G) and 42 (H). Arrows indicate enlarged nephron tubules and areas of fluid retention. Inflammation and atrophy are evident on days 14, 28 and 42.

### Hepatic CYP3A2 & CYP2C11 mRNA Expression over CKD Progression

CYP3A2 mRNA expression was minimally decreased on day 3, recovered on day 7, then declined substantially by day 14 (−83%, p<0.001) which persisted to day 42 (−99%, p<0.001) (Figure 2A). CYP2C11 mRNA expression was unchanged on day 3 but largely increased in the control group on day 7 leaving CKD rats well below normal (−76%, p<0.001) (Figure 2B). On day 14 (−84%, p<0.001), day 28 (−96%, p<0.001), and day 42 (−98%, p<0.001) the CKD CYP2C11 mRNA expression was decreased in comparison to control.

**Figure 2.**
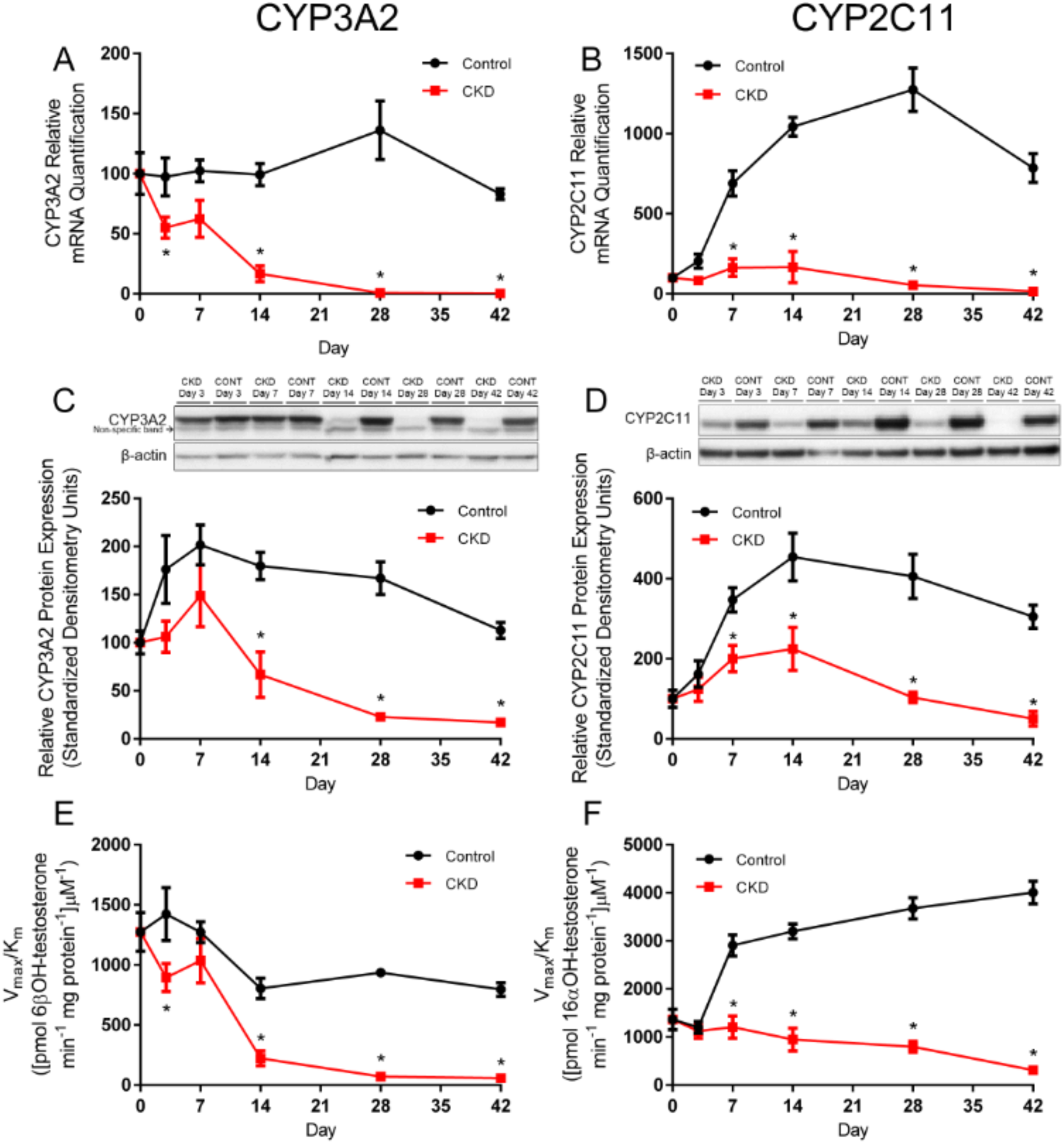
Relative mRNA expression, protein expression and enzymatic activity levels of CYP3A2 and CYP2C11. CYP3A2 (A) and CYP2C11 (B) mRNA expression and protein expression, CYP3A2 (C) and CYP2C11 (D), with representative western blots. Values were relative to β-actin represented as the mean ± SEM, normalized to control day 0 and arbitrarily defined as 100%. Enzymatic activity of CYP3A2 (D) and CYP2C11 (E) in control and CKD rats represented as the mean intrinsic clearance V_max_/K_m_ [(ml/min/mg protein)] of testosterone metabolite ± SEM. *p < 0.05 when compared to matching day control; n ≥ 6.

### Hepatic CYP3A2 & CYP2C11 Protein Expression over CKD Progression

Significant decreases in CYP3A2 protein expression were observed in CKD animals on day 14 (−63%, p<0.001), day 28 (−86%, p<0.001) and day 42 (−85%, p<0.01) (Figure 2C). CYP2C11 protein quantification also shows depletion in CKD but starting on day 7 (−42%, p<0.05) through to day 42 (−83%, p<0.001) (Figure 2D).

### Hepatic CYP3A2 & CYP2C11 Enzymatic Activity over CKD Progression

CYP3A2 intrinsic activity in CKD rats decreased on day 3, recovered on day 7 and fell again 3.6-fold by day 14, 13-fold by day 28 and nearly 14-fold lower than controls by day 42 (Figure 2E). The intrinsic activity of CYP2C11 showed a 4.6-fold difference between CKD and control as early as day 7 and up to 12.8-fold difference by day 42 (Figure 2F). Michaelis-Menten parameters are summarized (Table 1).

**Table 1.** CYP3A2 and CYP2C11 enzymatic activity over CKD progression. 6βOH-testosterone (CYP3A2) and 16αOH-testosterone (CYP2C11) production of liver microsomes measured following incubation with NADPH and testosterone. V_max_ values are in pmol/min/mg protein and Km values are µM. *p<0.05 compared to matching day control; n ≥ 6.

| **6βOH-testosterone (CYP3A2)** |  |  |  |  |  |  |
| --- | --- | --- | --- | --- | --- | --- |
| | $V_{\max}^a$ | | $K_m^a$ | | Intrinsic Clearance ($V_{\max}/K_m$)<sup>a</sup> | |
|  | Control | CKD | Control | CKD | Control | CKD |
| **Day 0** | 35938±2841 | 35938±2841 | 29.5±9.02 | 29.5±9.02 | 1275±393.87 | 1275±393.87 |
| **Day 3** | 56184±5640 | **26425±2097\*** | 40.1±14.46 | 29.8±9.13 | 1422±540.42 | **894±288.25\*** |
| **Day 7** | 49912±1806 | 42396±4911 | 40.2±5.218 | 42.7±17.48 | 1272±214.04 | 1036±489.05 |
| **Day 14** | 38223±2797 | **12804±1983\*** | 49.4±12.34 | 59.4±29.88 | 804±207.09 | **222±152.72\*** |
| **Day 28** | 61702±3069 | **10102±1289\*** | 66.6±10.43 | **141.6±44.74\*** | 936±86.36 | **71±21.50\*** |
| **Day 42** | 45706±2266 | **7849±1297\*** | 58.5±9.46 | **135.6±56.32\*** | 795±139.12 | **58±24.76\*** |
| **16αOH-testosterone (CYP2C11)** |  |  |  |  |  |  |
| | $V_{\max}^a$ | | $K_m^a$ | | Intrinsic Clearance ($V_{\max}/K_m$)<sup>a</sup> | |
|  | Control | CKD | Control | CKD | Control | CKD |
| **Day 0** | 17983±1210 | 17983±1210 | 14.2±4.56 | 14.2±4.56 | 1365±518.05 | 1365±518.05 |
| **Day 3** | 25892±2353 | 15453±1195 | 21.8±8.28 | 13.8±5.13 | 1202±298.73 | 1123±357.65 |
| **Day 7** | 41420±2488 | 27114±3431 | 14.3±4.09 | 23.5±12.17 | 2907±541.70 | **1206±607.94\*** |
| **Day 14** | 51824±2556 | **17521±2406\*** | 16.9±3.75 | 18.6±11.18 | 3199±377.15 | **948±523.91\*** |
| **Day 28** | 72358±2064 | **16236±1548\*** | 20.1±2.45 | 21.2±8.52 | 3679±548.84 | **797±273.19\*** |
| **Day 42** | 62571±3147 | **6658±513\*** | 16.2±3.71 | 22.6±7.21 | 4005±570.00 | **312±179.33\*** |

### Plasma & Liver Metabolomics

Untargeted metabolomics analysis was used to assess changes in metabolite composition. Principal component analysis (PCA) clearly separated CKD and control for both rat plasma and liver samples (Figure 3). Early disease stages are arbitrarily defined as day 3-14 and late stages days 28 and 42. R^2^ and Q^2^ parameters were used to accompany the interpretation of OPLS-DA plots (Table 2). Metabolites in rat plasma were well separated from control as early as day 3 when using RPLC (Figure 3A). Liver RPLC showed far less separation between control and CKD before day 28 (Figure 3B). The HILIC column showed separation back to day 7 except for poor Q^2^ values on day 14 in both plasma and liver samples (Figure 3 C-D).

**Table 2.**
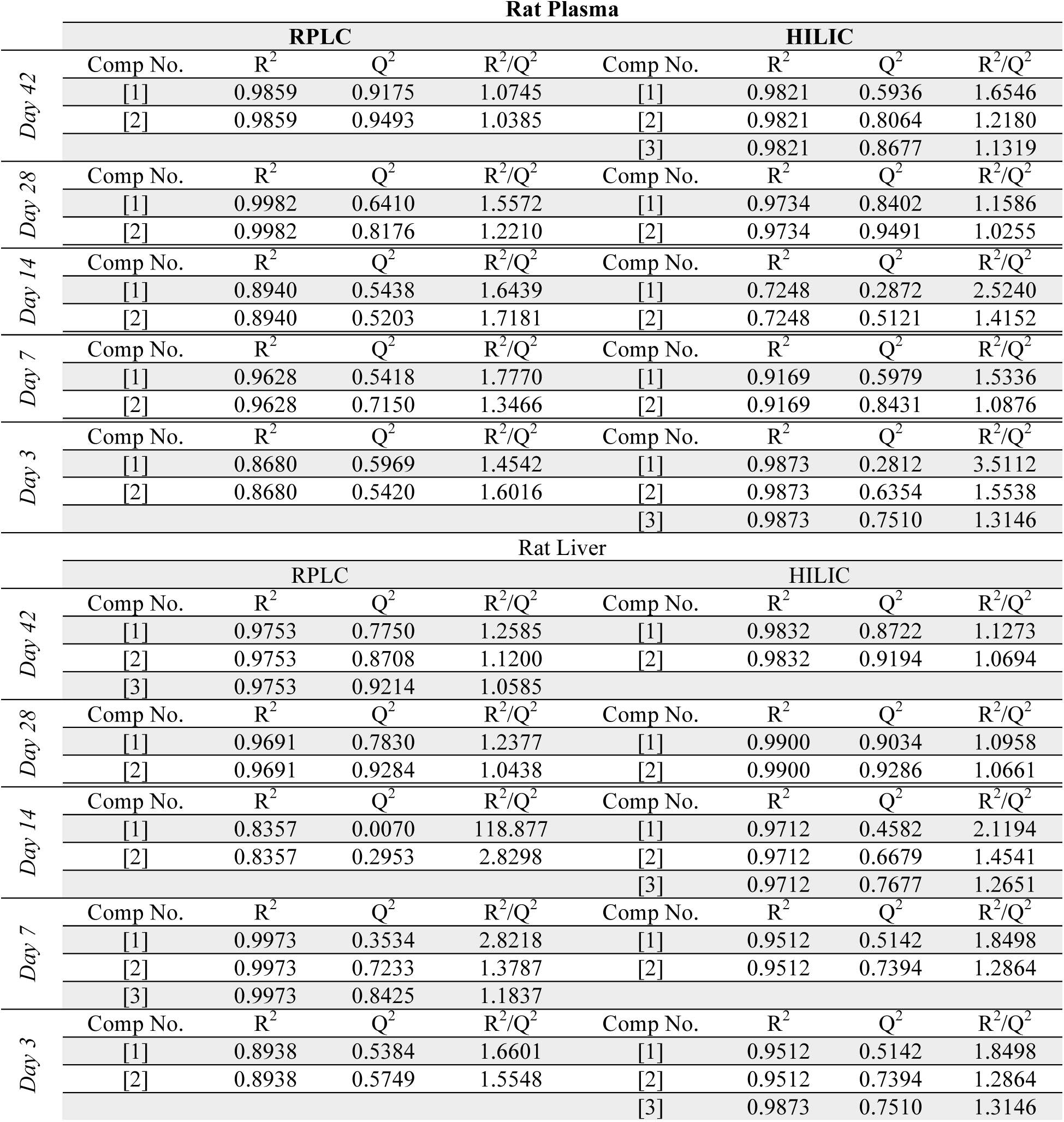
Multivariate OPLS-DA parameters R^2^ and Q^2^. R^2^ and Q^2^ values for plasma and liver metabolomics using RPLC and HILIC across all time points.

| Rat Plasma |  |  |  |  |  |  |  |  |
| --- | --- | --- | --- | --- | --- | --- | --- | --- |
|  | RPLC |  |  |  | HILIC |  |  |  |
| | Comp No. | $R^2$ | $Q^2$ | $R^2/Q^2$ | Comp No. | $R^2$ | $Q^2$ | $R^2/Q^2$ |
| Day 42 | [1] | 0.9859 | 0.9175 | 1.0745 | [1] | 0.9821 | 0.5936 | 1.6546 |
|  | [2] | 0.9859 | 0.9493 | 1.0385 | [2] | 0.9821 | 0.8064 | 1.2180 |
|  |  |  |  |  | [3] | 0.9821 | 0.8677 | 1.1319 |
| Day 28 | Comp No. | $R^2$ | $Q^2$ | $R^2/Q^2$ | Comp No. | $R^2$ | $Q^2$ | $R^2/Q^2$ |
|  | [1] | 0.9982 | 0.6410 | 1.5572 | [1] | 0.9734 | 0.8402 | 1.1586 |
|  | [2] | 0.9982 | 0.8176 | 1.2210 | [2] | 0.9734 | 0.9491 | 1.0255 |
| Day 14 | Comp No. | $R^2$ | $Q^2$ | $R^2/Q^2$ | Comp No. | $R^2$ | $Q^2$ | $R^2/Q^2$ |
|  | [1] | 0.8940 | 0.5438 | 1.6439 | [1] | 0.7248 | 0.2872 | 2.5240 |
|  | [2] | 0.8940 | 0.5203 | 1.7181 | [2] | 0.7248 | 0.5121 | 1.4152 |
| Day 7 | Comp No. | $R^2$ | $Q^2$ | $R^2/Q^2$ | Comp No. | $R^2$ | $Q^2$ | $R^2/Q^2$ |
|  | [1] | 0.9628 | 0.5418 | 1.7770 | [1] | 0.9169 | 0.5979 | 1.5336 |
|  | [2] | 0.9628 | 0.7150 | 1.3466 | [2] | 0.9169 | 0.8431 | 1.0876 |
| Day 3 | Comp No. | $R^2$ | $Q^2$ | $R^2/Q^2$ | Comp No. | $R^2$ | $Q^2$ | $R^2/Q^2$ |
|  | [1] | 0.8680 | 0.5969 | 1.4542 | [1] | 0.9873 | 0.2812 | 3.5112 |
|  | [2] | 0.8680 | 0.5420 | 1.6016 | [2] | 0.9873 | 0.6354 | 1.5538 |
|  |  |  |  |  | [3] | 0.9873 | 0.7510 | 1.3146 |
| Rat Liver |  |  |  |  |  |  |  |  |
|  | RPLC |  |  |  | HILIC |  |  |  |
| | Comp No. | $R^2$ | $Q^2$ | $R^2/Q^2$ | Comp No. | $R^2$ | $Q^2$ | $R^2/Q^2$ |
| Day 42 | [1] | 0.9753 | 0.7750 | 1.2585 | [1] | 0.9832 | 0.8722 | 1.1273 |
|  | [2] | 0.9753 | 0.8708 | 1.1200 | [2] | 0.9832 | 0.9194 | 1.0694 |
|  | [3] | 0.9753 | 0.9214 | 1.0585 |  |  |  |  |
| Day 28 | Comp No. | $R^2$ | $Q^2$ | $R^2/Q^2$ | Comp No. | $R^2$ | $Q^2$ | $R^2/Q^2$ |
|  | [1] | 0.9691 | 0.7830 | 1.2377 | [1] | 0.9900 | 0.9034 | 1.0958 |
|  | [2] | 0.9691 | 0.9284 | 1.0438 | [2] | 0.9900 | 0.9286 | 1.0661 |
| Day 14 | Comp No. | $R^2$ | $Q^2$ | $R^2/Q^2$ | Comp No. | $R^2$ | $Q^2$ | $R^2/Q^2$ |
|  | [1] | 0.8357 | 0.0070 | 118.877 | [1] | 0.9712 | 0.4582 | 2.1194 |
|  | [2] | 0.8357 | 0.2953 | 2.8298 | [2] | 0.9712 | 0.6679 | 1.4541 |
|  |  |  |  |  | [3] | 0.9712 | 0.7677 | 1.2651 |
| Day 7 | Comp No. | $R^2$ | $Q^2$ | $R^2/Q^2$ | Comp No. | $R^2$ | $Q^2$ | $R^2/Q^2$ |
|  | [1] | 0.9973 | 0.3534 | 2.8218 | [1] | 0.9512 | 0.5142 | 1.8498 |
|  | [2] | 0.9973 | 0.7233 | 1.3787 | [2] | 0.9512 | 0.7394 | 1.2864 |
|  | [3] | 0.9973 | 0.8425 | 1.1837 |  |  |  |  |
| Day 3 | Comp No. | $R^2$ | $Q^2$ | $R^2/Q^2$ | Comp No. | $R^2$ | $Q^2$ | $R^2/Q^2$ |
|  | [1] | 0.8938 | 0.5384 | 1.6601 | [1] | 0.9512 | 0.5142 | 1.8498 |
|  | [2] | 0.8938 | 0.5749 | 1.5548 | [2] | 0.9512 | 0.7394 | 1.2864 |
|  |  |  |  |  | [3] | 0.9873 | 0.7510 | 1.3146 |

**Figure 3.**
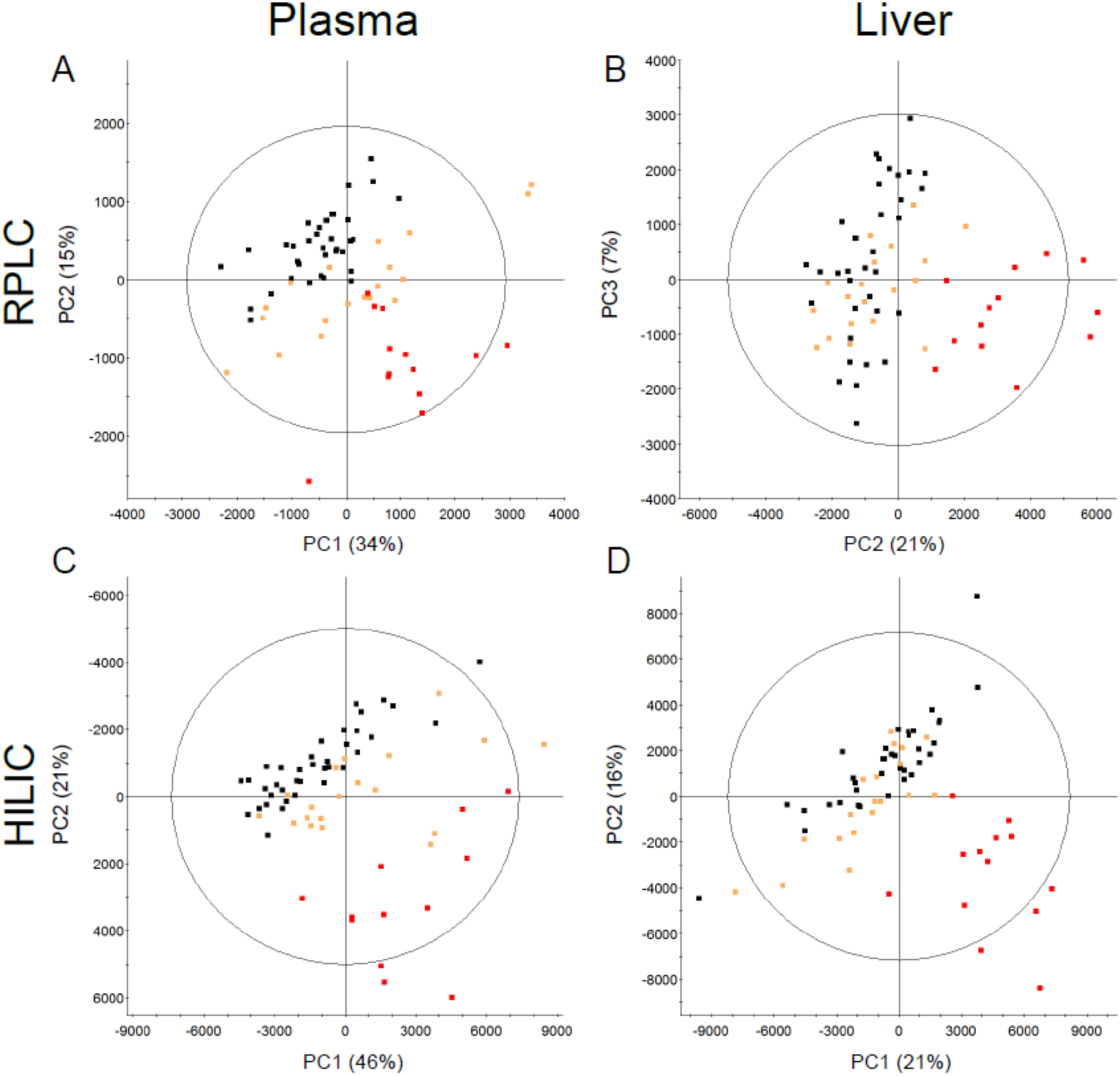
Unsupervised principal component analysis (PCA) plots of rat plasma (A) and liver (B) metabolome separated by RPLC. PCA of plasma (C) and liver (D) metabolome separated by HILIC. Each point is either control (▪), early stage CKD defined by day 3, 7 and 14 (▪), or late stage CKD defined by days 28 and 42 (▪). Each axis is either the first [1], second [2] or third [3] principal component showing the two components representing the largest variation between groups. Placement of each sample is determined by the metabolite composition within each sample and clustered samples share similar compositions. Data is centered and Pareto-scaled. Select rat samples were removed as outliers (A) no outliers, (B) a day 28 CKD sample, (C) a day 3 and day 42 control, and (D) a day 7 control sample.

### CYP Enzymes & Uremic Toxins

After satisfying multivariate and univariate analysis, metabolites were correlated to CYP mRNA, protein and enzymatic activity. There were 204 unique *m/z* ratios identified across all four runs that correlated with either CYP3A2 or CYP2C11 (Supplementary Table 4). Of these 204 *m/z* ratios, 9 metabolites were identified at identification level 1 using purchased standards. These metabolites include: allantoin, L-carnitine, creatinine, 2,8-dihydroxyadenine, equol-4/7-O-glucuronide, 4-ethylphenyl sulfate, indoxyl sulfate, pantothenic acid (vitamin B5) and phenyl sulfate (Table 3). Indoxyl sulfate, phenyl sulfate and 4-ethylphenyl sulfate had increased concentration (p<0.0001) on days 28 and 42 for both plasma and liver tissue (Figure 4).

**Figure 4.**
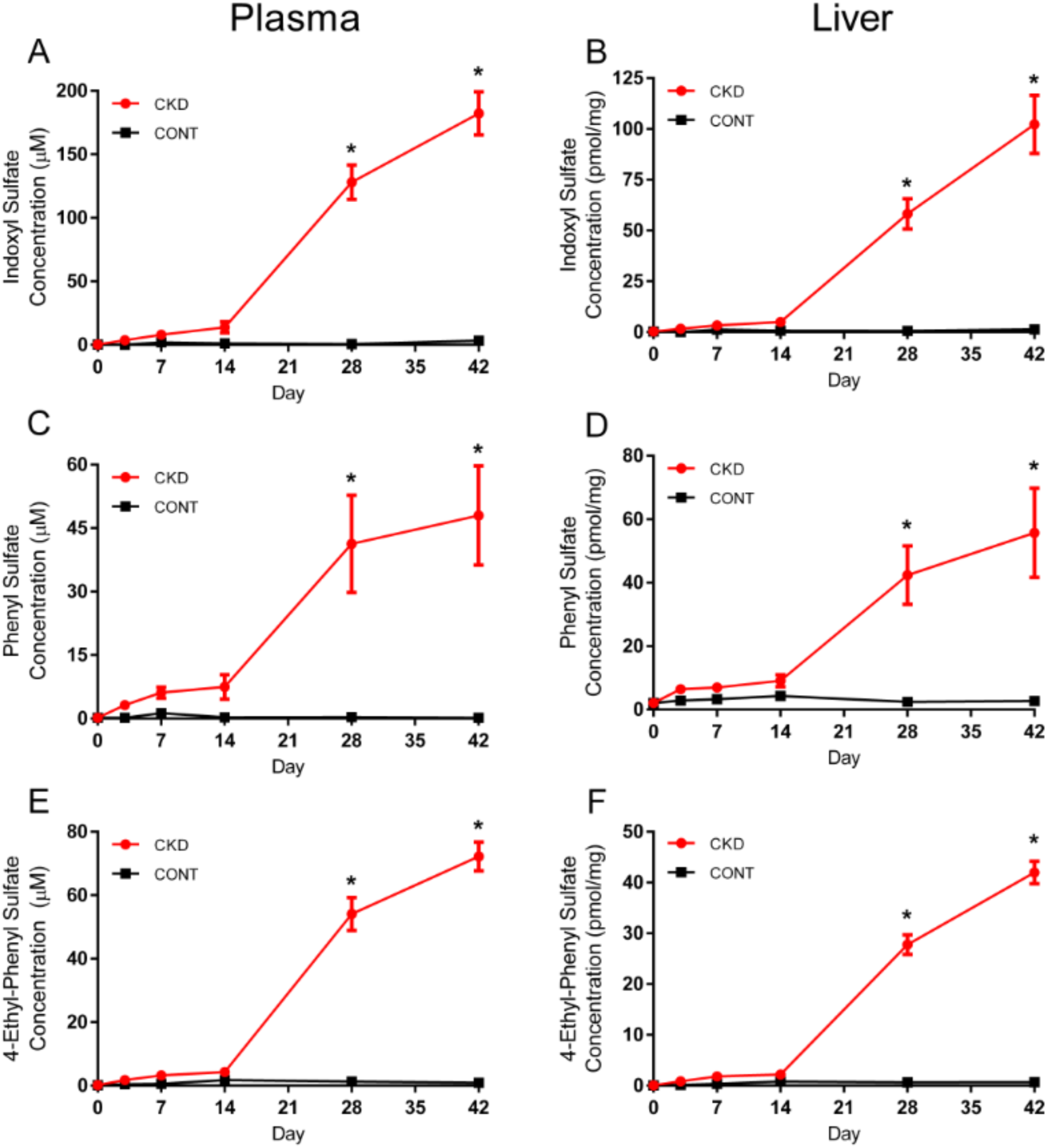
Quantitative analysis of metabolites indoxyl sulfate, phenyl sulfate and 4-ethylphenyl Sulfate. Plasma indoxyl sulfate (A), phenyl sulfate (C), 4-ethylphenyl sulfate (E) (µM) and liver indoxyl sulfate (B), phenyl sulfate (D) and 4-ethylphenyl sulfate (F) (pmol/mg liver tissue) concentrations obtained via untargeted metabolomics. Results are presented as mean ± SEM, \**p* < 0.0001 when compared to same day control; n≥6.

### Indoxyl Sulfate Downregulates Hepatic CYP3A4 Expression *in vitro*

To investigate the mechanism of CYP3A downregulation in CKD, human hepatoma Huh7 cells were treated with selected uremic toxins. As an initial proof-of-concept screen, concentrations of uremic toxins used were the highest reported in patients with CKD (41). A 24-hour indoxyl sulfate treatment decreased CYP3A4 mRNA expression by 70% in Huh7 cells (Figure 5A). Other individual uremic toxins did not affect CYP3A4 mRNA expression. Indoxyl sulfate is a protein bound uremic toxin; therefore, the concentration dependence of this effect was evaluated in the presence of 40 g/L HSA in the culture medium. Treatment with indoxyl sulfate for 48 hours produced a concentration-dependent decrease in CYP3A4 mRNA expression as the concentration was increased from those measured in healthy controls to concentrations in patients with CKD (IC_50_ = 113.0 ± 3.5 µM) (Figure 5B). A significant decrease in the steady-state levels of CYP3A4 mRNA was demonstrated when Huh7 cells were treated with ≥ 300 µM indoxyl sulfate in the presence of 40 g/L HSA supplemented media (Figure 5B, *p* < 0.05). Huh7 cells treated with indoxyl sulfate in the uremic range resulted in a 21% to 95% decrease in CYP3A4 mRNA expression. Cell viability was unaffected by indoxyl sulfate in media containing HSA after 48 hour treatment at clinically relevant concentrations (Figure 5C).

**Figure 5.**
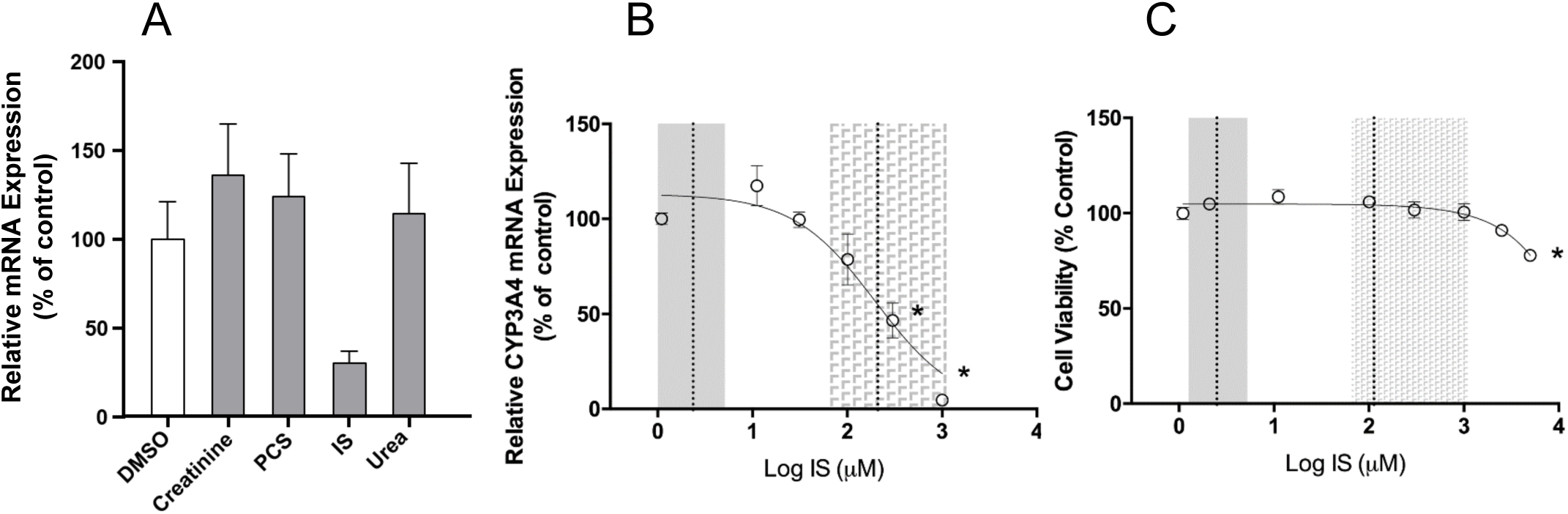
A) The effect of select uremic toxins on CYP3A4 expression in Huh7 cells. B) Concentration dependent effect of indoxyl sulfate on CYP3A4 expression in Huh7 cells. C) Cell viability in the presence of increasing indoxyl sulfate concentrations. Results are represented as mean ± SEM, \**p* < 0.05; n ≥ 3.

### Caecal Microbiota

To understand if the gut microbiota was changing in parallel with metabolite changes, next-generation Illumina sequencing was used to assess the bacterial composition of the caecum (Supplementary Table 2). Exploratory PCA ordination showed that the caecum samples had high intrinsic biological variation, but separated by time, regardless of disease state (Figure 6A). A multivariate analysis found that CKD and control separated into distinct groups at day 28 and 42 (day 28: R^2^ = 0.97; Q^2^ =0.71 and day 42: R^2^ = 0.98; Q^2^ =0.70) (Figure 6B). Effect size and overlap of each OTU relative abundance was tabulated and assessed for trends. Only two bacterial OTUs changed between control and CKD on two or more consecutive days with respect to effect size and overlap. The first OTU was from the phylum Firmicutes and genus Turicibacter and was significantly higher in CKD rats compared to control animals on days 14, 28 and 42 (Figure 7A) with an increasing trend associated with disease progression. The second OTU from phylum Bacteroidetes and genus Parabacteroides showed a significant decrease in control rats over time (Figure 7B).

**Figure 6.**
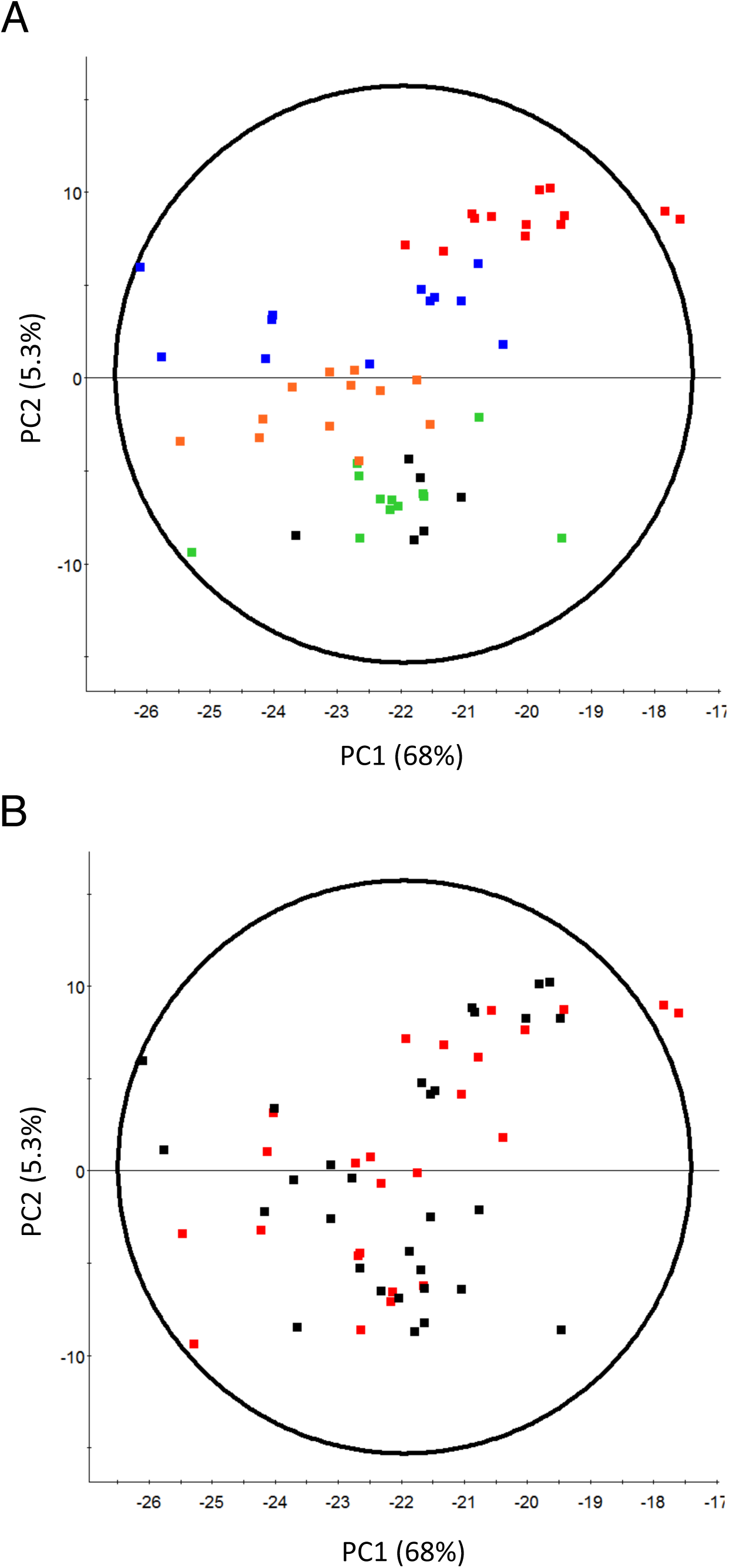
Unsupervised principal component analysis (PCA) of control and CKD rat caecum bacterial sequences coloured by (A) day 0 (▪) 3 (▪), 14 (▪), 28 (▪) and 42 (▪) or by (B) treatment, CKD (▪) or control (▪). Data is centered without scaling.

**Figure 7.**
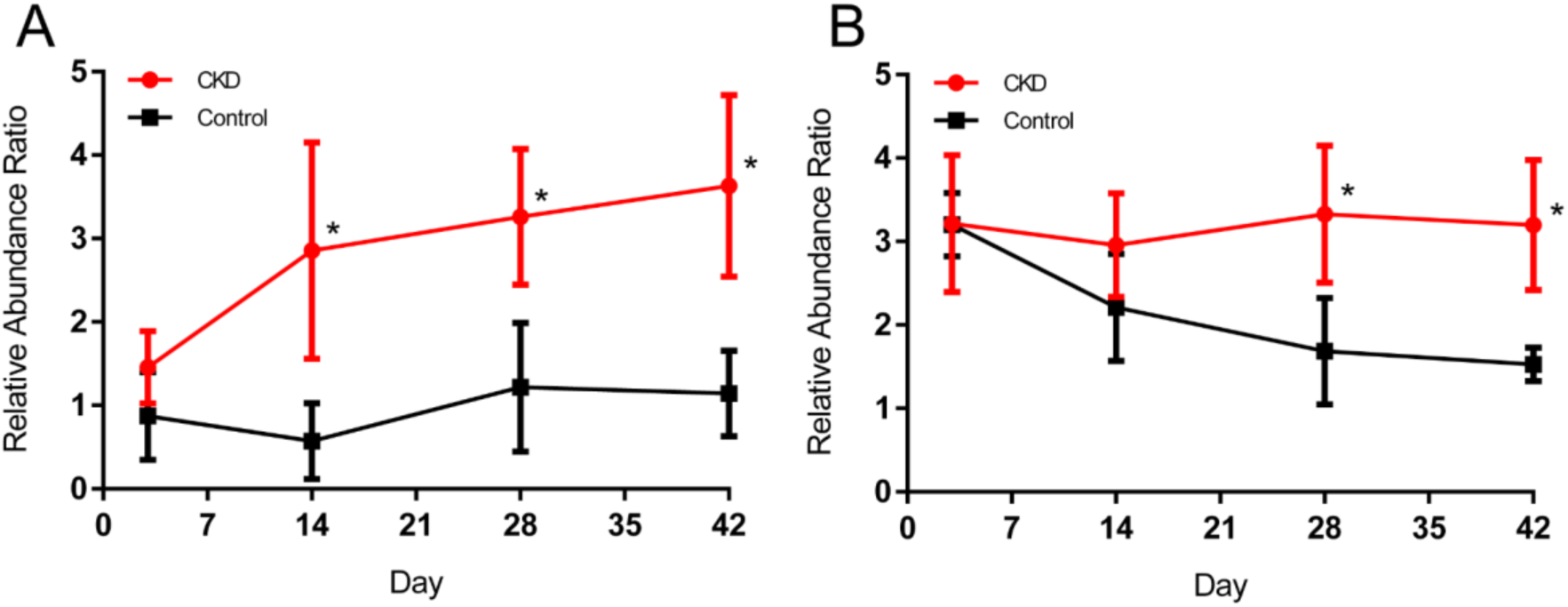
Mean relative abundance of genus Turicibacter (A) and genus Parabacteroides (B) displayed as the CLR-transformed values of the OTUs ± 95% confidence interval using R v3.2.3 package ALDEx2 v1.2.0. *Absolute effect size ≥ 1.5 and overlap < 6.5% compared to same day control; n ≥ 6.

## Discussion

The pharmacokinetics of many drugs are unpredictably altered in CKD, making these patients susceptible to adverse drug events. Hepatic CYPs play a crucial role in nonrenal drug clearance and alterations in these CYPs may contribute to pharmacokinetic changes observed in CKD. CYP downregulation has been associated with decreased renal clearance and consequent retention of uremic toxins in animal models of CKD. Mechanistic studies suggest the involvement of various pathways from pre-transcriptional regulation to direct inhibition by uremic toxins, inflammatory factors, and hormones (9–13, 42, 43). Uremic toxins are also suggested to change the relative abundance of gut bacteria to favor uremic-toxin producing microbes and create a state of dysbiosis in CKD patients (44). Changes in metabolite or toxin production as a consequence of the altered microbiota may exacerbate CKD progression and potentially CYP downregulation. However, the pathophysiological factors of uremia and dysbiosis have yet to be evaluated temporally. In this manuscript, uremia and dysbiosis were characterized over CKD progression to identify potential causes of CYP downregulation.

CYP3A2 and CYP2C11 were both downregulated by CKD with respect to mRNA expression, protein expression, and enzymatic activity as previously observed (45). These findings suggest these CYPs were initially influenced at the transcriptional level, consequently leading to altered protein expression and enzyme activity. Expression of CYP3A2 in control rats was stable throughout the study while expression started to decrease on day 14 in CKD rats. These observations suggest the removal, inhibition, or downregulation of a constitutive factor required for expression, potentially mediated by the increase in uremic toxins (46). In contrast, control rats exhibited an increase in CYP2C11 expression as early as day 7 but this increase was not evident in CKD rats. Increased CYP2C11 expression over time has been described in healthy male juvenile rats, where it was suspected to reflect increased testosterone levels during puberty (46). Furthermore, CKD has been associated with hypogonadism and testosterone deficiency (47). CYP2C11 is also influenced by alterations in the normally cyclic levels of growth hormone (GH) where continuous GH release or loss of GH production will both downregulate CYP2C11 (48).

The differing trends between CYP3A2 and CYP2C11 may be attributed to nuclear receptor differences. CYP2C11 is less dependent on hepatocyte nuclear factor 4 alpha induction, and CKD-induced receptor binding inhibition is less extensive for CYP2C11 than it is CYP3A2 (17, 49). Alternatively, it is also possible that removal or inhibition of shared nuclear receptor PXR or reduced receptor binding of RNA polymerase II is affecting both enzymes but in different manners depending on substrate availability (9, 17).

Plasma samples showed greater metabolomics separation in earlier stages of disease (days 3-14) than liver samples. This suggests CKD first inflicts a uremic environment in the plasma before infiltrating the liver. Uremic changes also overlapped with the early changes in CYP3A2 and CYP2C11, supporting the hypothesis that uremic toxins are involved with CYP regulation. Metabolites from each metabolomics run were subjected to correlation analysis with CYP3A2 or CYP2C11 mRNA, protein or enzymatic activity levels. Of the 204 *m/z* features retrieved, 8 of the 9 identified at level 1 classification were increased with CKD progression [allantoin, creatinine, 2,8-dihydroxyadenine, pantothenic acid (vitamin B_5_), indoxyl sulfate, phenyl sulfate, equol-4/7- O-glucuronide and 4-ethylphenyl sulfate]. L-carnitine was the only level 1 metabolite that showed a positive correlation with CYP downregulation, decreasing over CKD progression.

Gut derived uremic toxins that are potentially involved in CYP downregulation from this study include indoxyl sulfate, phenyl sulfate, 4-ethylphenyl sulfate, equol-4/7-O-glucournide and products of L-carnitine metabolism. Indoxyl sulfate and phenyl sulfate are two highly retained gut-derived uremic toxins (50, 51) both found in CKD patients and animal models (18, 52). Indoxyl sulfate and phenyl sulfate have been associated with altered drug metabolism both through transcriptional regulation (42), and indoxyl sulfate as a direct inhibitor of CYP activity (13). Thus, the identification and observed increase in concentration of indoxyl sulfate and phenyl sulfate in this study support their previously described roles in modifying CYP regulation in CKD (Figure 4). Further, our in vitro studies using Huh7 human hepatoma cells showed indoxyl sulfate decreases CYP3A4 expression in a concentration dependent manner. Interestingly, of the metabolites found by correlation to CYP3A2 or CYP2C11, the five metabolites that are associated with CYP downregulation in the literature are all gut-derived uremic toxins.

Indoxyl sulfate, phenyl sulfate and 4-ethylphenyl sulfate concentration all increase after day 28 when changes are simultaneously observed in the gut microbiota that was phylogenetically analyzed using 16S sequencing. This lends support to the idea that uremia may be driving the change in gut microbial abundance through a damaged gut wall (Figure 8) (44, 53). The late and pronounced increase in gut-derived uremic toxins also suggests dysbiosis contributes to the exacerbation of uremia, likely adding to the uremic milieu by increasing the number of bacteria capable of uremic toxin production (44). Multivariate analysis showed the microbiota was most significantly influenced by time (i.e. age) and secondarily by disease state. This correlation suggests that the microbiota changes caused by CKD induction are less profound than age-associated bacterial changes. Additionally, in comparison to the metabolomic PCA, microbial clustering with respect to disease state was poor. This may indicate that the uremic environment in the plasma and liver are altered prior to the onset of dysbiosis. Bacterial families significantly changed on days 28 and 42 contained strains capable of producing at least one of the following genes: urease, tryptophanase, phosphotransbutyrylase or butyrate kinase, although the bacteria were inconsistently characteristic of either control or CKD rats (54). The sole bacterial genus significantly changed due to disease state prior to day 14 was of the order Clostridiales, in accordance with the findings of Barrios and colleagues who sequenced the gut microbiota of 855 people and correlated bacteria from the order Clostridiales with early renal decline (55).

**Figure 8.**
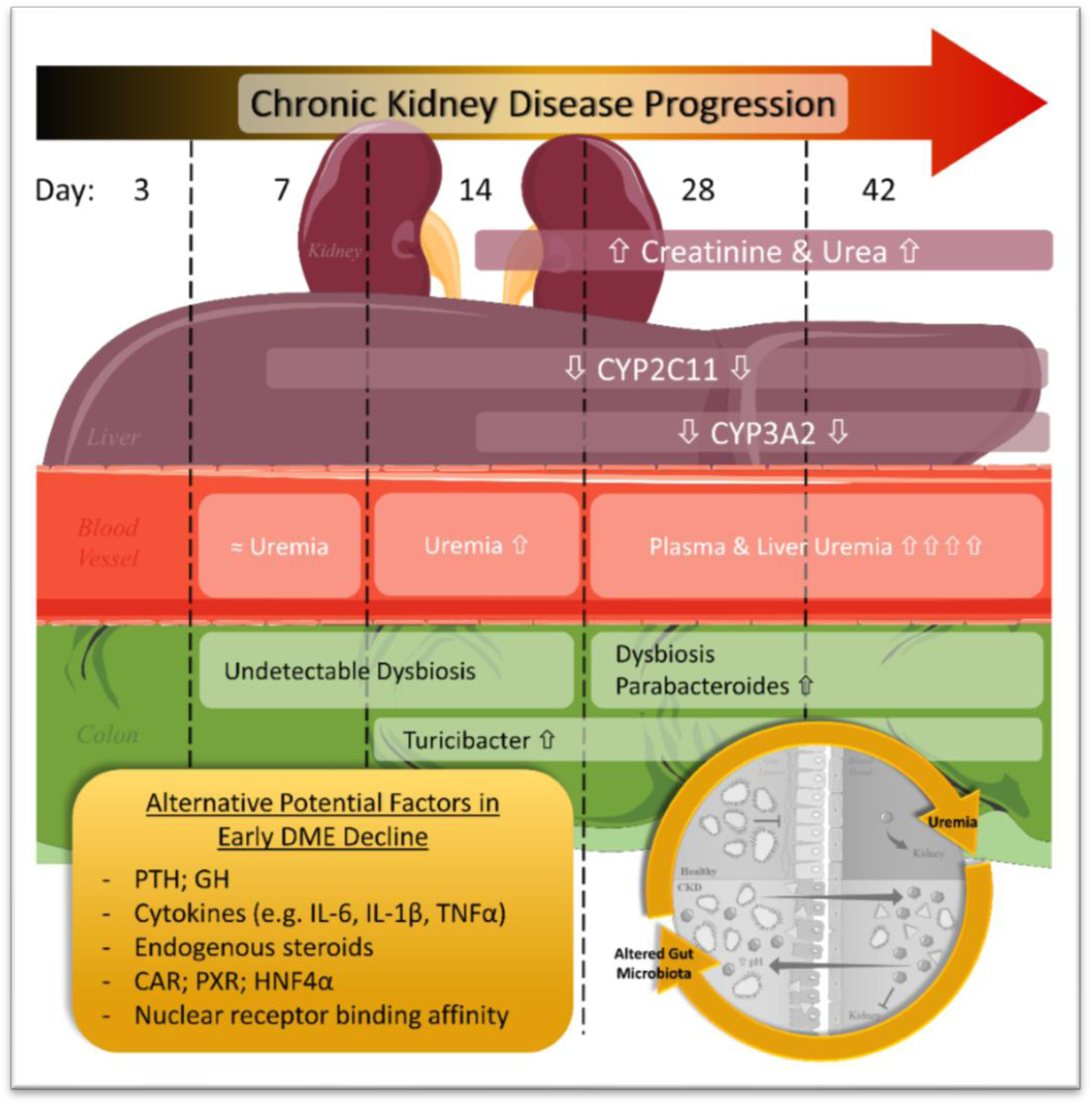
Temporal associations of uremia and dysbiosis with CYP3A2 and CYP2C11 expression over CKD progression. Days (3 through 42) refer to rat study time points carried out in this thesis. CKD was characterized by urea and creatinine beginning on day 14 and correlating with the decrease in CYP3A2 expression. CYP2C11 expression decreased as early as day 7. Although plasma uremia may be involved as early as day 7 indicated by untargeted multivariate analysis, quantified uremic toxins were significantly increased only on days 28 and 42. Similarly, gut bacterial dysbiosis was detectable on days 28 and 42 supporting the hypothesis of a positive-feedback cycle involving uremia and the gut microbiota. This study suggests there are likely other factors influencing CYPs in early stages of CKD. Images were modified from Servier Medical Art (http://www.servier.co.uk/medical-art-gallery).

Although interesting to examine the temporal relationship for OTUs differing due to CKD, only two bacterial genera, *Turicibacter* and *Parabacteroides*, were significant on two or more consecutive days, best correlating with CYP trends. *Turicibacter* was the most consistently changed bacteria, changing as early as day 14 through to day 42 with an increasing trend as CKD progressed. Identifying the genus *Turicibacter* in CKD animals is a novel finding. *Turicibacter* are gram-positive, strictly anaerobic, rod-shaped bacteria of which very little is known. *Turicibacter* has been identified in the blood of febrile patients with acute appendicitis (56) and associated with pouchitis – a complication of proctocolectomy – in ulcerative colitis patients (57). A fecal microbiota transplant from healthy human into colons of germ-free rats also identifies *Turicibacter sp* (58). Only 4 strains, within the *sanguinis* species, have been published to date: MOL361 (56), PC909 (59), ZCY83 (60), H121 (61). A BLASTn search of our *Turicibacter* sequence matched the MOL361 species with 100% identity (NR_028816.1). Assuming all rats were exposed to *Turicibacter* for the study duration, our findings suggest CKD animals are more susceptible to gut colonization by *Turicibacter*.

The BLASTn results for the *Parabacteroides* genus OTU suggested 99% sequence identity to two stains of the species *distasonis*: strain ATCC 8503 (NR_074376.1) and JCM 5825 (NR_041342.1). In 2006, *Bacteroides distasonis* was reclassified as *Parabacteroides distasonis* and thus, all subsequent information pertains to either classification (62). *Parabacteroides* is a gram-negative, anaerobic, non-spore-forming genus. *P. distasonis* is classified in the KEGG pathway database as an opportunistic pathogen capable of anaerobic infection (63). Analysis of bacteria capable of generating phenol and indole compounds found *P. distasonis* proficient at producing p-cresol (64) and indoxyl sulfate (65). In general, it seems *P. distasonis* is potentially both harmful and beneficial depending on translocation and abundance. Our results show a unique trend where CKD rats have a stable level of *Parabacteroides* and controls slowly reduce the abundance of this genus after 28 days. Given the multitude of associations with disease, *Parabacteroides* may be taking advantage of the dysbiotic state in CKD when it is normally removed in controls by other healthy bacteria as a part of the progression in age-associated microbial changes.

In conclusion, global plasma and liver alterations of the metabolome over disease progression provide support for uremic toxins playing a role in CYP downregulation. Alternatively, the early detection of CYP downregulation and late surge of gut-derived uremic toxin concentrations suggest other factors are involved in CYP regulation in early stages of CKD (Figure 8). A temporal association was established between severe CKD, caecal dysbiosis and increase in gut-derived uremic toxins indoxyl sulfate, phenyl sulfate and 4-ethylphenyl sulfate. This association supports the positive-feedback loop of uremia and dysbiosis suspected to drive severe CKD (Figure 8).

## Abbreviations

CKD: Chronic Kidney Disease;
CYP: Cytochrome P-450;
ESI: Electrospray ionization;
GFR: Glomerular Filtration Rate;
GH: Growth Hormone;
HILIC: Hydrophilic Interaction Liquid Chromatography;
IS: Indoxyl Sulfate;
MS: Mass Spectrometry;
OTU: Operational Taxonomic Unit;
OPLS-DA: Orthogonal Partial Least Squares Discriminant Analysis;
PTH: Parathyroid Hormone;
PS: Phenyl Sulfate;
PCA: Principal Component Analysis;
QTof/MS: Quadrupole Time-of-flight Mass Spectrometer;
RPLC: Reverse Phase Liquid Chromatography;
UPLC: Ultra-Performance Liquid Chromatography;
VIP: Variable Importance in Projection

