## Supplementary Information for "The Impact of Uremia and Intestinal Dysbiosis on Hepatic Drug Metabolism in a Rat Model of Progressive Chronic Kidney Disease" (from another version of this preprint: version 1, 2019-01-27)

**Supplementary Table 1.** Table of barcoded primers used for Illumina sequencing.

| Plate_well | Left Primer | Right Primer | Left Barcode | Right Barcode | Sample ID |
|---|---|---|---|---|---|
| Plate2_A1 | V4L5 | V5R1 | ccttggaa | ccaaggtt | DNA-Blank-1 |
| Plate2_B1 | V4L5 | V5R21 | ccttggaa | cacagtgt | 445-Ceacum-11-03-15 |
| Plate2_C1 | V4L6 | V5R1 | ttggaacc | ccaaggtt | 465-Ceacum-12-14-15 |
| Plate2_D1 | V4L6 | V5R21 | ttggaacc | cacagtgt | 415-Ceacum-10-06-15 |
| Plate2_E1 | V4L7 | V5R1 | ggaacctt | ccaaggtt | 455-Ceacum-12-10-15 |
| Plate2_F1 | V4L7 | V5R21 | ggaacctt | cacagtgt | 444-Stool-11-03-15 |
| Plate2_G1 | V4L8 | V5R1 | aaccttgg | ccaaggtt | 447-Stool-09-22-15 |
| Plate2_H1 | V4L8 | V5R21 | aaccttgg | cacagtgt | 478-Stool-01-18-16 |
| Plate2_A2 | V4L5 | V5R2 | ccttggaa | aaggttcc | 484-Stool-01-18-16 |
| Plate2_B2 | V4L5 | V5R22 | ccttggaa | agagtctc | 398-Ceacum-09-22-15 |
| Plate2_C2 | V4L6 | V5R2 | ttggaacc | aaggttcc | 469-Ceacum-12-14-15 |
| Plate2_D2 | V4L6 | V5R22 | ttggaacc | agagtctc | 440-Stool-11-03-15 |
| Plate2_E2 | V4L7 | V5R2 | ggaacctt | aaggttcc | 419-Stool-10-06-15 |
| Plate2_F2 | V4L7 | V5R22 | ggaacctt | agagtctc | 394-Stool-09-22-15 |
| Plate2_G2 | V4L8 | V5R2 | aaccttgg | aaggttcc | 416-Stool-10-06-15 |
| Plate2_H2 | V4L8 | V5R22 | aaccttgg | agagtctc | -- |
| Plate2_A3 | V4L5 | V5R3 | ccttggaa | ggttccaa | 464-Stool-12-14-15 |
| Plate2_B3 | V4L5 | V5R23 | ccttggaa | tctcagag | 474-Stool-12-14-15 |
| Plate2_C3 | V4L6 | V5R3 | ttggaacc | ggttccaa | 459-Stool-12-10-15 |
| Plate2_D3 | V4L6 | V5R23 | ttggaacc | tctcagag | 401-Ceacum-09-25-15 |
| Plate2_E3 | V4L7 | V5R3 | ggaacctt | ggttccaa | 437-Stool-09-25-15 |
| Plate2_F3 | V4L7 | V5R23 | ggaacctt | tctcagag | 483-Stool-01-18-16 |
| Plate2_G3 | V4L8 | V5R3 | aaccttgg | ggttccaa | 469-Stool-12-14-15 |
| Plate2_H3 | V4L8 | V5R23 | aaccttgg | tctcagag | -- |
| Plate2_A4 | V4L5 | V5R4 | ccttggaa | ttccaagg | 403-Stool-09-25-15 |
| Plate2_B4 | V4L5 | V5R24 | ccttggaa | gagacaca | 408-Stool-09-25-15 |
| Plate2_C4 | V4L6 | V5R4 | ttggaacc | ttccaagg | 417-Ceacum-10-06-15 |
| Plate2_D4 | V4L6 | V5R24 | ttggaacc | gagacaca | 446-Stool-11-03-15 |
| Plate2_E4 | V4L7 | V5R4 | ggaacctt | ttccaagg | 429-Stool-10-20-15 |
| Plate2_F4 | V4L7 | V5R24 | ggaacctt | gagacaca | 462-Ceacum-12-14-15 |
| Plate2_G4 | V4L8 | V5R4 | aaccttgg | ttccaagg | 445-Stool-11-03-15 |
| Plate2_H4 | V4L8 | V5R24 | aaccttgg | gagacaca | -- |
| Plate2_A5 | V4L5 | V5R5 | ccttggaa | ccttggaa | 452-Stool-12-07-15 |
| Plate2_B5 | V4L5 | V5R25 | ccttggaa | ctactacc | 434-Stool-10-20-15 |
| Plate2_C5 | V4L6 | V5R5 | ttggaacc | ccttggaa | 405-Ceacum-09-25-15 |
| Plate2_D5 | V4L6 | V5R25 | ttggaacc | ctactacc | 419-Ceacum-10-06-15 |
| Plate2_E5 | V4L7 | V5R5 | ggaacctt | ccttggaa | 475-Ceacum-01-18-16 |
| Plate2_F5 | V4L7 | V5R25 | ggaacctt | ctactacc | 436-Stool-09-25-15 |
| Plate2_G5 | V4L8 | V5R5 | aaccttgg | ccttggaa | 436-Ceacum-11-03-15 |
| Plate2_H5 | V4L8 | V5R25 | aaccttgg | ctactacc | -- |
| Plate2_A6 | V4L5 | V5R6 | ccttggaa | ttggaacc | 471-Ceacum-12-14-15 |
| Plate2_B6 | V4L5 | V5R26 | ccttggaa | gatgatgg | 438-Stool-11-03-15 |

| | | | | | |
|---|---|---|---|---|---|
| Plate2_C6 | V4L6 | V5R6 | ttggaacc | ttggaacc | 460-Stool-12-10-15 |
| Plate2_D6 | V4L6 | V5R26 | ttggaacc | gatgatgg | 395-Ceacum-09-22-15 |
| Plate2_E6 | V4L7 | V5R6 | ggaacctt | ttggaacc | 477-Stool-01-18-16 |
| Plate2_F6 | V4L7 | V5R26 | ggaacctt | gatgatgg | 441-Stool-09-22-15 |
| Plate2_G6 | V4L8 | V5R6 | aaccttgg | ttggaacc | 448-Ceacum-12-07-15 |
| Plate2_H6 | V4L8 | V5R26 | aaccttgg | gatgatgg | -- |
| Plate2_A7 | V4L5 | V5R7 | ccttggaa | ggaacctt | 445-Stool-09-22-15 |
| Plate2_B7 | V4L5 | V5R27 | ccttggaa | tcgtcgtt | 460-Ceacum-12-10-15 |
| Plate2_C7 | V4L6 | V5R7 | ttggaacc | ggaacctt | 473-Stool-12-14-15 |
| Plate2_D7 | V4L6 | V5R27 | ttggaacc | tcgtcgtt | 432-Stool-10-20-15 |
| Plate2_E7 | V4L7 | V5R7 | ggaacctt | ggaacctt | 439-Stool-09-22-15 |
| Plate2_F7 | V4L7 | V5R27 | ggaacctt | tcgtcgtt | 459-Ceacum-12-10-15 |
| Plate2_G7 | V4L8 | V5R7 | aaccttgg | ggaacctt | 454-Ceacum-12-10-15 |
| Plate2_H7 | V4L8 | V5R27 | aaccttgg | tcgtcgtt | -- |
| Plate2_A8 | V4L5 | V5R8 | ccttggaa | aaccttgg | 463-Ceacum-12-14-15 |
| Plate2_B8 | V4L5 | V5R28 | ccttggaa | agcagcaa | 399-Ceacum-09-25-15 |
| Plate2_C8 | V4L6 | V5R8 | ttggaacc | aaccttgg | 438-Stool-09-25-15 |
| Plate2_D8 | V4L6 | V5R28 | ttggaacc | agcagcaa | 405-Stool-09-25-15 |
| Plate2_E8 | V4L7 | V5R8 | ggaacctt | aaccttgg | 476-Stool-01-18-16 |
| Plate2_F8 | V4L7 | V5R28 | ggaacctt | agcagcaa | 482-Ceacum-01-18-16 |
| Plate2_G8 | V4L8 | V5R8 | aaccttgg | aaccttgg | 457-Stool-12-10-15 |
| Plate2_H8 | V4L8 | V5R28 | aaccttgg | agcagcaa | -- |
| Plate2_A9 | V4L5 | V5R17 | ccttggaa | ggttaacc | 446-Stool-10-20-15 |
| Plate2_B9 | V4L5 | V5R29 | ccttggaa | ctaccсta | 447-Stool-11-03-15 |
| Plate2_C9 | V4L6 | V5R17 | ttggaacc | ggttaacc | 409-Stool-09-25-15 |
| Plate2_D9 | V4L6 | V5R29 | ttggaacc | ctacccta | 443-Stool-09-22-15 |
| Plate2_E9 | V4L7 | V5R17 | ggaacctt | ggttaacc | 410-Ceacum-09-25-15 |
| Plate2_F9 | V4L7 | V5R29 | ggaacctt | ctacccta | 444-Stool-09-25-15 |
| Plate2_G9 | V4L8 | V5R17 | aaccttgg | ggttaacc | 484-Ceacum-01-18-16 |
| Plate2_H9 | V4L8 | V5R29 | aaccttgg | ctacccta | -- |
| Plate2_A10 | V4L5 | V5R18 | ccttggaa | ttaaccgg | 479-Stool-01-18-16 |
| Plate2_B10 | V4L5 | V5R30 | ccttggaa | gatgggat | 426-Stool-10-20-15 |
| Plate2_C10 | V4L6 | V5R18 | ttggaacc | ttaaccgg | 397-Stool-09-22-15 |
| Plate2_D10 | V4L6 | V5R30 | ttggaacc | gatgggat | 443-Stool-11-03-15 |
| Plate2_E10 | V4L7 | V5R18 | ggaacctt | ttaaccgg | 486-Stool-01-18-16 |
| Plate2_F10 | V4L7 | V5R30 | ggaacctt | gatgggat | 461-Ceacum-12-10-15 |
| Plate2_G10 | V4L8 | V5R18 | aaccttgg | ttaaccgg | 435-Stool-09-25-15 |
| Plate2_H10 | V4L8 | V5R30 | aaccttgg | gatgggat | -- |
| Plate2_A11 | V4L5 | V5R19 | ccttggaa | aaccggtt | 435-Ceacum-11-03-15 |
| Plate2_B11 | V4L5 | V5R31 | ccttggaa | cgtttcgt | 458-Ceacum-12-10-15 |
| Plate2_C11 | V4L6 | V5R19 | ttggaacc | aaccggtt | 439-Stool-10-20-15 |
| Plate2_D11 | V4L6 | V5R31 | ttggaacc | cgtttcgt | 394-Ceacum-09-22-15 |
| Plate2_E11 | V4L7 | V5R19 | ggaacctt | aaccggtt | 441-Stool-11-03-15 |
| Plate2_F11 | V4L7 | V5R31 | ggaacctt | cgtttcgt | 435-Stool-11-03-15 |
| Plate2_G11 | V4L8 | V5R19 | aaccttgg | aaccggtt | 439-Ceacum-11-03-15 |

| | | | | | |
|---|---|---|---|---|---|
| Plate2_H11 | V4L8 | V5R31 | aaccttgg | cgtttcgt | -- |
| Plate2_A12 | V4L5 | V5R20 | ccttggaa | ccggttaa | 422-Ceacum-10-06-15 |
| Plate2_B12 | V4L5 | V5R32 | ccttggaa | gcaaagca | 451-Ceacum-12-07-15 |
| Plate2_C12 | V4L6 | V5R20 | ttggaacc | ccggttaa | 425-Ceacum-10-20-15 |
| Plate2_D12 | V4L6 | V5R32 | ttggaacc | gcaaagca | 480-Stool-01-18-16 |
| Plate2_E12 | V4L7 | V5R20 | ggaacctt | ccggttaa | 401-Stool-09-25-15 |
| Plate2_F12 | V4L7 | V5R32 | ggaacctt | gcaaagca | 430-Ceacum-10-20-15 |
| Plate2_G12 | V4L8 | V5R20 | aaccttgg | ccggttaa | 461-Stool-12-10-15 |
| Plate2_H12 | V4L8 | V5R32 | aaccttgg | gcaaagca | PCR-Blank-1 |
| Plate5_A1 | V4L25 | V5R1 | ctactacc | ccaaggtt | 395-Stool-09-22-15 |
| Plate5_B1 | V4L25 | V5R21 | ctactacc | cacagtgt | 404-Ceacum-09-25-15 |
| Plate5_C1 | V4L26 | V5R1 | gatgatgg | ccaaggtt | 468-Stool-12-14-15 |
| Plate5_D1 | V4L26 | V5R21 | gatgatgg | cacagtgt | 468-Ceacum-12-14-15 |
| Plate5_E1 | V4L27 | V5R1 | tcgtcgtt | ccaaggtt | 440-Stool-09-22-15 |
| Plate5_F1 | V4L27 | V5R21 | tcgtcgtt | cacagtgt | 425-Stool-10-20-15 |
| Plate5_G1 | V4L28 | V5R1 | agcagcaa | ccaaggtt | 447-Stool-09-25-15 |
| Plate5_H1 | V4L28 | V5R21 | agcagcaa | cacagtgt | -X- |
| Plate5_A2 | V4L25 | V5R2 | ctactacc | aaggttcc | DNA-Blank-2 |
| Plate5_B2 | V4L25 | V5R22 | ctactacc | agagtctc | 448-Stool-12-07-15 |
| Plate5_C2 | V4L26 | V5R2 | gatgatgg | aaggttcc | 472-Stool-12-14-15 |
| Plate5_D2 | V4L26 | V5R22 | gatgatgg | agagtctc | 443-Stool-10-20-15 |
| Plate5_E2 | V4L27 | V5R2 | tcgtcgtt | aaggttcc | 449-Ceacum-12-07-15 |
| Plate5_F2 | V4L27 | V5R22 | tcgtcgtt | agagtctc | 411-Stool-10-06-15 |
| Plate5_G2 | V4L28 | V5R2 | agcagcaa | aaggttcc | 424-Stool-10-20-15 |
| Plate5_H2 | V4L28 | V5R22 | agcagcaa | agagtctc | -X- |
| Plate5_A3 | V4L25 | V5R3 | ctactacc | ggttccaa | 445-Stool-09-25-15 |
| Plate5_B3 | V4L25 | V5R23 | ctactacc | tctcagag | 464-Ceacum-12-14-15 |
| Plate5_C3 | V4L26 | V5R3 | gatgatgg | ggttccaa | 420-Ceacum-10-06-15 |
| Plate5_D3 | V4L26 | V5R23 | gatgatgg | tctcagag | 428-Stool-10-20-15 |
| Plate5_E3 | V4L27 | V5R3 | tcgtcgtt | ggttccaa | 480-Ceacum-01-18-16 |
| Plate5_F3 | V4L27 | V5R23 | tcgtcgtt | tctcagag | 483-Ceacum-01-18-16 |
| Plate5_G3 | V4L28 | V5R3 | agcagcaa | ggttccaa | 443-Stool-10-06-15 |
| Plate5_H3 | V4L28 | V5R23 | agcagcaa | tctcagag | -X- |
| Plate5_A4 | V4L25 | V5R4 | ctactacc | ttccaagg | 417-Stool-10-06-15 |
| Plate5_B4 | V4L25 | V5R24 | ctactacc | gagacaca | 418-Stool-10-06-15 |
| Plate5_C4 | V4L26 | V5R4 | gatgatgg | ttccaagg | 473-Ceacum-12-14-15 |
| Plate5_D4 | V4L26 | V5R24 | gatgatgg | gagacaca | 400-Stool-09-25-15 |
| Plate5_E4 | V4L27 | V5R4 | tcgtcgtt | ttccaagg | 438-Ceacum-11-03-15 |
| Plate5_F4 | V4L27 | V5R24 | tcgtcgtt | gagacaca | 447-Stool-10-20-15 |
| Plate5_G4 | V4L28 | V5R4 | agcagcaa | ttccaagg | 440-Stool-10-06-15 |
| Plate5_H4 | V4L28 | V5R24 | agcagcaa | gagacaca | -X- |
| Plate5_A5 | V4L25 | V5R5 | ctactacc | ccttggaa | 406-Ceacum-09-25-15 |
| Plate5_B5 | V4L25 | V5R25 | ctactacc | ctactacc | 458-Stool-12-10-15 |
| Plate5_C5 | V4L26 | V5R5 | gatgatgg | ccttggaa | 416-Ceacum-10-06-15 |
| Plate5_D5 | V4L26 | V5R25 | gatgatgg | ctactacc | 445-Stool-10-20-15 |

| | | | | | |
|---|---|---|---|---|---|
| Plate5_E5 | V4L27 | V5R5 | tcgtcgtt | ccttggaa | 393-Stool-09-22-15 |
| Plate5_F5 | V4L27 | V5R25 | tcgtcgtt | ctactacc | 431-Stool-10-20-15 |
| Plate5_G5 | V4L28 | V5R5 | agcagcaa | ccttggaa | -X- |
| Plate5_H5 | V4L28 | V5R25 | agcagcaa | ctactacc | -X- |
| Plate5_A6 | V4L25 | V5R6 | ctactacc | ttggaacc | 439-Stool-11-03-15 |
| Plate5_B6 | V4L25 | V5R26 | ctactacc | gatgatgg | 462-Stool-12-14-15 |
| Plate5_C6 | V4L26 | V5R6 | gatgatgg | ttggaacc | 408-Ceacum-09-25-15 |
| Plate5_D6 | V4L26 | V5R26 | gatgatgg | gatgatgg | 398-Stool-09-22-15 |
| Plate5_E6 | V4L27 | V5R6 | tcgtcgtt | ttggaacc | 426-Ceacum-10-20-15 |
| Plate5_F6 | V4L27 | V5R26 | tcgtcgtt | gatgatgg | 421-Ceacum-10-06-15 |
| Plate5_G6 | V4L28 | V5R6 | agcagcaa | ttggaacc | -X- |
| Plate5_H6 | V4L28 | V5R26 | agcagcaa | gatgatgg | -X- |
| Plate5_A7 | V4L25 | V5R7 | ctactacc | ggaacctt | 412-Ceacum-10-06-15 |
| Plate5_B7 | V4L25 | V5R27 | ctactacc | tcgtcgtt | 453-Stool-12-07-15 |
| Plate5_C7 | V4L26 | V5R7 | gatgatgg | ggaacctt | 456-Stool-12-10-15 |
| Plate5_D7 | V4L26 | V5R27 | gatgatgg | tcgtcgtt | 403-Ceacum-09-25-15 |
| Plate5_E7 | V4L27 | V5R7 | tcgtcgtt | ggaacctt | 432-Ceacum-10-20-15 |
| Plate5_F7 | V4L27 | V5R27 | tcgtcgtt | tcgtcgtt | 471-Stool-12-14-15 |
| Plate5_G7 | V4L28 | V5R7 | agcagcaa | ggaacctt | -X- |
| Plate5_H7 | V4L28 | V5R27 | agcagcaa | tcgtcgtt | -X- |
| Plate5_A8 | V4L25 | V5R8 | ctactacc | aaccttgg | 393-Ceacum-09-22-15 |
| Plate5_B8 | V4L25 | V5R28 | ctactacc | agcagcaa | 409-Ceacum-09-25-15 |
| Plate5_C8 | V4L26 | V5R8 | gatgatgg | aaccttgg | 467-Stool-12-14-15 |
| Plate5_D8 | V4L26 | V5R28 | gatgatgg | agcagcaa | 438-Stool-10-20-15 |
| Plate5_E8 | V4L27 | V5R8 | tcgtcgtt | aaccttgg | 428-Ceacum-10-20-15 |
| Plate5_F8 | V4L27 | V5R28 | tcgtcgtt | agcagcaa | 453-Ceacum-12-07-15 |
| Plate5_G8 | V4L28 | V5R8 | agcagcaa | aaccttgg | -X- |
| Plate5_H8 | V4L28 | V5R28 | agcagcaa | agcagcaa | -X- |
| Plate5_A9 | V4L25 | V5R17 | ctactacc | ggttaacc | 449-Stool-12-07-15 |
| Plate5_B9 | V4L25 | V5R29 | ctactacc | ctacccta | 440-Ceacum-11-03-15 |
| Plate5_C9 | V4L26 | V5R17 | gatgatgg | ggttaacc | 472-Ceacum-12-14-15 |
| Plate5_D9 | V4L26 | V5R29 | gatgatgg | ctacccta | 435-Stool-10-20-15 |
| Plate5_E9 | V4L27 | V5R17 | tcgtcgtt | ggttaacc | 421-Stool-10-06-15 |
| Plate5_F9 | V4L27 | V5R29 | tcgtcgtt | ctacccta | 437-Stool-10-06-15 |
| Plate5_G9 | V4L28 | V5R17 | agcagcaa | ggttaacc | -X- |
| Plate5_H9 | V4L28 | V5R29 | agcagcaa | ctacccta | -X- |
| Plate5_A10 | V4L25 | V5R18 | ctactacc | ttaaccgg | 441-Stool-10-20-15 |
| Plate5_B10 | V4L25 | V5R30 | ctactacc | gatgggat | 429-Ceacum-10-20-15 |
| Plate5_C10 | V4L26 | V5R18 | gatgatgg | ttaaccgg | 439-Stool-09-25-15 |
| Plate5_D10 | V4L26 | V5R30 | gatgatgg | gatgggat | 437-Stool-10-20-15 |
| Plate5_E10 | V4L27 | V5R18 | tcgtcgtt | ttaaccgg | 450-Ceacum-12-07-15 |
| Plate5_F10 | V4L27 | V5R30 | tcgtcgtt | gatgggat | 407-Stool-09-25-15 |
| Plate5_G10 | V4L28 | V5R18 | agcagcaa | ttaaccgg | -X- |
| Plate5_H10 | V4L28 | V5R30 | agcagcaa | gatgggat | -X- |
| Plate5_A11 | V4L25 | V5R19 | ctactacc | aaccggtt | 406-Stool-09-25-15 |

| | | | | | |
|---|---|---|---|---|---|
| Plate5_B11 | V4L25 | V5R31 | ctactacc | cgtttcgt | 410-Stool-09-25-15 |
| Plate5_C11 | V4L26 | V5R19 | gatgatgg | aaccggtt | 487-Ceacum-01-18-16 |
| Plate5_D11 | V4L26 | V5R31 | gatgatgg | cgtttcgt | 434-Ceacum-10-20-15 |
| Plate5_E11 | V4L27 | V5R19 | tcgtcgtt | aaccggtt | 463-Stool-12-14-15 |
| Plate5_F11 | V4L27 | V5R31 | tcgtcgtt | cgtttcgt | 470-Ceacum-12-14-15 |
| Plate5_G11 | V4L28 | V5R19 | agcagcaa | aaccggtt | -X- |
| Plate5_H11 | V4L28 | V5R31 | agcagcaa | cgtttcgt | -X- |
| Plate5_A12 | V4L25 | V5R20 | ctactacc | ccggttaa | 414-Stool-10-06-15 |
| Plate5_B12 | V4L25 | V5R32 | ctactacc | gcaaagca | 447-Ceacum-11-03-15 |
| Plate5_C12 | V4L26 | V5R20 | gatgatgg | ccggttaa | 441-Stool-10-06-15 |
| Plate5_D12 | V4L26 | V5R32 | gatgatgg | gcaaagca | 442-Stool-10-20-15 |
| Plate5_E12 | V4L27 | V5R20 | tcgtcgtt | ccggttaa | 474-Ceacum-12-14-15 |
| Plate5_F12 | V4L27 | V5R32 | tcgtcgtt | gcaaagca | 418-Ceacum-10-06-15 |
| Plate5_G12 | V4L28 | V5R20 | agcagcaa | ccggttaa | -X- |
| Plate5_H12 | V4L28 | V5R32 | agcagcaa | gcaaagca | PCR-Blank-2 |
| Plate6_A1 | V4L29 | V5R1 | ctacccta | ccaaggtt | DNA-Blank-3 |
| Plate6_B1 | V4L29 | V5R21 | ctacccta | cacagtgt | 454-Stool-12-10-15 |
| Plate6_C1 | V4L30 | V5R1 | gatgggat | ccaaggtt | 396-Ceacum-09-22-15 |
| Plate6_D1 | V4L30 | V5R21 | gatgggat | cacagtgt | 442-Ceacum-11-03-15 |
| Plate6_E1 | V4L31 | V5R1 | cgtttcgt | ccaaggtt | 481-Ceacum-01-18-16 |
| Plate6_F1 | V4L31 | V5R21 | cgtttcgt | cacagtgt | 487-Stool-01-18-16 |
| Plate6_G1 | V4L32 | V5R1 | gcaaagca | ccaaggtt | 444-Stool-10-20-15 |
| Plate6_H1 | V4L32 | V5R21 | gcaaagca | cacagtgt | -X- |
| Plate6_A2 | V4L29 | V5R2 | ctacccta | aaggttcc | 436-Stool-11-03-15 |
| Plate6_B2 | V4L29 | V5R22 | ctacccta | agagtctc | 486-Ceacum-01-18-15 |
| Plate6_C2 | V4L30 | V5R2 | gatgggat | aaggttcc | 450-Stool-12-07-15 |
| Plate6_D2 | V4L30 | V5R22 | gatgggat | agagtctc | 402-Stool-09-25-15 |
| Plate6_E2 | V4L31 | V5R2 | cgtttcgt | aaggttcc | 440-Stool-09-25-15 |
| Plate6_F2 | V4L31 | V5R22 | cgtttcgt | agagtctc | 435-Stool-10-06-15 |
| Plate6_G2 | V4L32 | V5R2 | gcaaagca | aaggttcc | 431-Ceacum-10-20-15 |
| Plate6_H2 | V4L32 | V5R22 | gcaaagca | agagtctc | -X- |
| Plate6_A3 | V4L29 | V5R3 | ctacccta | ggttccaa | 430-Stool-10-20-15 |
| Plate6_B3 | V4L29 | V5R23 | ctacccta | tctcagag | 476-Ceacum-01-18-16 |
| Plate6_C3 | V4L30 | V5R3 | gatgggat | ggttccaa | 444-Stool-10-06-15 |
| Plate6_D3 | V4L30 | V5R23 | gatgggat | tctcagag | 413-Ceacum-10-06-15 |
| Plate6_E3 | V4L31 | V5R3 | cgtttcgt | ggttccaa | 411-Ceacum-10-06-15 |
| Plate6_F3 | V4L31 | V5R23 | cgtttcgt | tctcagag | 442-Stool-09-22-15 |
| Plate6_G3 | V4L32 | V5R3 | gcaaagca | ggttccaa | 441-Stool-09-25-15 |
| Plate6_H3 | V4L32 | V5R23 | gcaaagca | tctcagag | -X- |
| Plate6_A4 | V4L29 | V5R4 | ctacccta | ttccaagg | 437-Stool-11-03-15 |
| Plate6_B4 | V4L29 | V5R24 | ctacccta | gagacaca | 451-Stool-12-07-15 |
| Plate6_C4 | V4L30 | V5R4 | gatgggat | ttccaagg | 436-Stool-09-22-15 |
| Plate6_D4 | V4L30 | V5R24 | gatgggat | gagacaca | 452-Ceacum-12-07-15 |
| Plate6_E4 | V4L31 | V5R4 | cgtttcgt | ttccaagg | 422-Stool-10-06-15 |
| Plate6_F4 | V4L31 | V5R24 | cgtttcgt | gagacaca | 437-Ceacum-11-03-15 |

| | | | | | |
|---|---|---|---|---|---|
| Plate6_G4 | V4L32 | V5R4 | gcaaagca | ttccaagg | 445-Stool-10-06-15 |
| Plate6_H4 | V4L32 | V5R24 | gcaaagca | gagacaca | -X- |
| Plate6_A5 | V4L29 | V5R5 | ctacccta | ccttggaa | 407-Ceacum-09-25-15 |
| Plate6_B5 | V4L29 | V5R25 | ctacccta | ctactacc | 455-Stool-12-10-15 |
| Plate6_C5 | V4L30 | V5R5 | gatgggat | ccttggaa | 465-Stool-12-14-15 |
| Plate6_D5 | V4L30 | V5R25 | gatgggat | ctactacc | 400-Ceacum-09-25-15 |
| Plate6_E5 | V4L31 | V5R5 | cgtttcgt | ccttggaa | 446-Stool-09-25-15 |
| Plate6_F5 | V4L31 | V5R25 | cgtttcgt | ctactacc | 470-Stool-12-14-15 |
| Plate6_G5 | V4L32 | V5R5 | gcaaagca | ccttggaa | 457-Ceacum-12-10-15 |
| Plate6_H5 | V4L32 | V5R25 | gcaaagca | ctactacc | -X- |
| Plate6_A6 | V4L29 | V5R6 | ctacccta | ttggaacc | 466-Ceacum-12-14-15 |
| Plate6_B6 | V4L29 | V5R26 | ctacccta | gatgatgg | 396-Stool-09-22-15 |
| Plate6_C6 | V4L30 | V5R6 | gatgggat | ttggaacc | 427-Ceacum-10-20-15 |
| Plate6_D6 | V4L30 | V5R26 | gatgggat | gatgatgg | 482-Stool-01-18-16 |
| Plate6_E6 | V4L31 | V5R6 | cgtttcgt | ttggaacc | 443-Ceacum-11-03-15 |
| Plate6_F6 | V4L31 | V5R26 | cgtttcgt | gatgatgg | 475-Stool-01-18-16 |
| Plate6_G6 | V4L32 | V5R6 | gcaaagca | ttggaacc | -X- |
| Plate6_H6 | V4L32 | V5R26 | gcaaagca | gatgatgg | -X- |
| Plate6_A7 | V4L29 | V5R7 | ctacccta | ggaacctt | 485-Ceacum-01-18-15 |
| Plate6_B7 | V4L29 | V5R27 | ctacccta | tcgtcgtt | 442-Stool-10-06-15 |
| Plate6_C7 | V4L30 | V5R7 | gatgggat | ggaacctt | 399-Stool-09-25-15 |
| Plate6_D7 | V4L30 | V5R27 | gatgggat | tcgtcgtt | 414-Ceacum-10-06-15 |
| Plate6_E7 | V4L31 | V5R7 | cgtttcgt | ggaacctt | 423-Ceacum-10-20-15 |
| Plate6_F7 | V4L31 | V5R27 | cgtttcgt | tcgtcgtt | 438-Stool-10-06-15 |
| Plate6_G7 | V4L32 | V5R7 | gcaaagca | ggaacctt | -X- |
| Plate6_H7 | V4L32 | V5R27 | gcaaagca | tcgtcgtt | -X- |
| Plate6_A8 | V4L29 | V5R8 | ctacccta | aaccttgg | 413-Stool-10-06-15 |
| Plate6_B8 | V4L29 | V5R28 | ctacccta | agcagcaa | 444-Ceacum-11-03-15 |
| Plate6_C8 | V4L30 | V5R8 | gatgggat | aaccttgg | 437-Stool-09-22-15 |
| Plate6_D8 | V4L30 | V5R28 | gatgggat | agcagcaa | 481-Stool-01-18-16 |
| Plate6_E8 | V4L31 | V5R8 | cgtttcgt | aaccttgg | 402-Ceacum-09-25-15 |
| Plate6_F8 | V4L31 | V5R28 | cgtttcgt | agcagcaa | 420-Stool-10-06-15 |
| Plate6_G8 | V4L32 | V5R8 | gcaaagca | aaccttgg | -X- |
| Plate6_H8 | V4L32 | V5R28 | gcaaagca | agcagcaa | -X- |
| Plate6_A9 | V4L29 | V5R17 | ctacccta | ggttaacc | 479-Ceacum-01-18-15 |
| Plate6_B9 | V4L29 | V5R29 | ctacccta | ctacccta | 442-Stool-11-03-15 |
| Plate6_C9 | V4L30 | V5R17 | gatgggat | ggttaacc | 412-Stool-10-06-15 |
| Plate6_D9 | V4L30 | V5R29 | gatgggat | ctacccta | 424-Ceacum-10-20-15 |
| Plate6_E9 | V4L31 | V5R17 | cgtttcgt | ggttaacc | 478-Ceacum-01-18-15 |
| Plate6_F9 | V4L31 | V5R29 | cgtttcgt | ctacccta | 404-Stool-9-25-15 |
| Plate6_G9 | V4L32 | V5R17 | gcaaagca | ggttaacc | -X- |
| Plate6_H9 | V4L32 | V5R29 | gcaaagca | ctacccta | -X- |
| Plate6_A10 | V4L29 | V5R18 | ctacccta | ttaaccgg | 477-Ceacum-01-18-15 |
| Plate6_B10 | V4L29 | V5R30 | ctacccta | gatgggat | 423-Stool-10-20-15 |
| Plate6_C10 | V4L30 | V5R18 | gatgggat | ttaaccgg | 456-Ceacum-12-10-15 |

| | | | | | |
|---|---|---|---|---|---|
| Plate6_D10 | V4L30 | V5R30 | gatgggat | gatgggat | 447-Stool-10-06-15 |
| Plate6_E10 | V4L31 | V5R18 | cgtttcgt | ttaaccgg | 439-Stool-10-06-15 |
| Plate6_F10 | V4L31 | V5R30 | cgtttcgt | gatgggat | 440-Stool-10-20-15 |
| Plate6_G10 | V4L32 | V5R18 | gcaaagca | ttaaccgg | -X- |
| Plate6_H10 | V4L32 | V5R30 | gcaaagca | gatgggat | -X- |
| Plate6_A11 | V4L29 | V5R19 | ctacccta | aaccggtt | 446-Stool-10-06-15 |
| Plate6_B11 | V4L29 | V5R31 | ctacccta | cgtttcgt | 433-Ceacum-10-20-15 |
| Plate6_C11 | V4L30 | V5R19 | gatgggat | aaccggtt | 438-Stool-09-22-15 |
| Plate6_D11 | V4L30 | V5R31 | gatgggat | cgtttcgt | 397-Ceacum-09-22-15 |
| Plate6_E11 | V4L31 | V5R19 | cgtttcgt | aaccggtt | 441-Ceacum-11-03-15 |
| Plate6_F11 | V4L31 | V5R31 | cgtttcgt | cgtttcgt | 467-Ceacum-12-14-15 |
| Plate6_G11 | V4L32 | V5R19 | gcaaagca | aaccggtt | -X- |
| Plate6_H11 | V4L32 | V5R31 | gcaaagca | cgtttcgt | -X- |
| Plate6_A12 | V4L29 | V5R20 | ctacccta | ccggttaa | 485-Stool-01-18-16 |
| Plate6_B12 | V4L29 | V5R32 | ctacccta | gcaaagca | 446-Ceacum-11-03-15 |
| Plate6_C12 | V4L30 | V5R20 | gatgggat | ccggttaa | 415-Stool-10-06-15 |
| Plate6_D12 | V4L30 | V5R32 | gatgggat | gcaaagca | 436-Stool-10-20-15 |
| Plate6_E12 | V4L31 | V5R20 | cgtttcgt | ccggttaa | 427-Stool-10-20-15 |
| Plate6_F12 | V4L31 | V5R32 | cgtttcgt | gcaaagca | 466-Stool-12-14-15 |
| Plate6_G12 | V4L32 | V5R20 | gcaaagca | ccggttaa | -X- |
| Plate6_H12 | V4L32 | V5R32 | gcaaagca | gcaaagca | PCR-Blank-3 |

**Supplementary Table 2.** Table of operational taxonomic units identified from Illumina sequencing. Please refer to attached excel file.

**Supplementary Table 3.** Weight in grams of control and CKD rats over 42 days.

| | | Control | CKD |
|---|---|---|---|
| *Weight (g)* | Day 0 | 206.5 ± 1.4 (n=41) | 207.1 ± 1.8 (n=36) |
| | Day 3 | 188.6 ± 1.1 (n=41) | 186.1 ± 1.5 (n=36) |
| | Day 7 | 201.0 ± 1.5 (n=24) | 202.4 ± 1.9 (n=26) |
| | Day 14 | 227.3 ± 2.0 (n=18) | 227.9 ± 3.3 (n=19) |
| | Day 28 | 257.3 ± 2.6 (n=12) | 258.3 ± 5.0 (n=13) |
| | Day 42 | 274.5 ± 8.4 (n=6) | 254.7 ± 12.7 (n=7) |

**Supplementary Table 4.** Table of 204 features found by untargeted mass spectrometry from CKD and control rat plasma and liver samples and RPLC or HILIC chromatography. Metabolites satisfying univariate analysis and Spearman correlation to same-sample CYP3A2 or CYP2C11 mRNA, protein or enzyme activity data. Spearman correlation coefficients (r value) are listed. 2-way independent ANOVA was conducted via MetaboAnalyst v3.0 using FDR<0.05 to correct for multiple comparisons and satisfaction required p<0.05 across both *Time* and *Disease*. Multivariate analysis required VIP > 0.8 and 0.4 < p(corr)[1] < -0.4 indicating adequate separation by OPLA-DA and S-plot. Mass error was obtained using the 4th decimal place *m/z* from the Human Metabolome Database (HMDB). Italicized suspected metabolites refer to a group of plausible metabolites of similar structure.

| ID | **Mass Error (ppm) from HMDB** | **Identity Level** | **Suspected Identity (Adduct)** | **Mass (m/z)** | **$t_R$ (min)** | **Biological Matrix** | **Column** | **ESI Mode** | **CYP3A2 Correlation** | | **CYP2C11 Correlation** | | **Satisfies Multivariate Analysis** |
|---|---|---|---|---|---|---|---|---|---|---|---|---|---|
| | | | | | | | | | **r value** | **mRNA/Protein /Activity** | **r value** | **mRNA/Protein /Activity** | |
| 1 | 1 | 1 | 2,8-Dihydroxyadenine | 168.0515 | 4.56 | Plasma | HILIC | Pos | **-0.7336** | mRNA | -0.6864 | Activity | FALSE |
| 2 | 3 | 1 | 4-ethyl-phenyl sulfate | 201.0221 | 1.06 | Liver | HILIC | Neg | -0.6689 | Activity | 0 | 0 | TRUE |
| 3 | 3 | 1 | 4-ethyl-phenyl sulfate | 201.0221 | 2.10 | Plasma | RPLC | Neg | -0.6813 | mRNA/**Activity** | 0 | 0 | TRUE |
| 4 | 3 | 1 | 4-ethyl-phenyl sulfate | 201.0221 | 2.13 | Liver | RPLC | Neg | **-0.7387** | mRNA/**Activity** | 0 | 0 | TRUE |
| 5 | 5 | 1 | Allantoin (M-H) | 157.0359 | 0.56 | Liver | RPLC | Neg | **-0.7284** | mRNA | **-0.7919** | mRNA/**Activity** | FALSE |
| 6 | 5 | 1 | Allantoin (M-H) | 157.0360 | 0.57 | Plasma | RPLC | Neg | **-0.7391** | mRNA | **-0.7921** | mRNA/**Activity** | FALSE |
| 7 | 2 | 1 | Allantoin (M+Na) | 181.0328 | 0.57 | Plasma | RPLC | Pos | **-0.7254** | mRNA | **-0.7073** | Activity | FALSE |
| 8 | 0 | 1 | Creatinine (M+H) | 114.0662 | 3.24 | Plasma | HILIC | Pos | **-0.7029** | mRNA/**Activity** | -0.6587 | Activity | TRUE |
| 9 | 2 | 1 | Equol 4/7-O-glucuronide | 417.1182 | 1.85 | Liver | RPLC | Neg | **-0.7265** | mRNA/**Protein** | **-0.805** | mRNA/Protein/ **Activity** | TRUE |
| 10 | 3 | 1 | Indoxyl sulfate | 212.0016 | 1.65 | Liver | RPLC | Neg | **-0.7201** | **mRNA**/Activity | 0 | 0 | TRUE |
| 11 | 3 | 1 | Indoxyl sulfate | 212.0017 | 1.65 | Plasma | RPLC | Neg | **-0.7072** | **mRNA**/Activity | 0 | 0 | TRUE |
| 12 | 1 | 1 | L-Carnitine | 162.1123 | 0.53 | Plasma | RPLC | Pos | 0.6694 | mRNA | **0.8657** | mRNA/Protein/ **Activity** | TRUE |
| 13 | 0 | 1 | Pantothenic acid (vitamin B5) | 220.1179 | 2.23 | Plasma | HILIC | Pos | 0 | 0 | **-0.7155** | mRNA/**Activity** | TRUE |
| 14 | 4 | 1 | Phenyl sulfate | 172.9907 | 1.38 | Plasma | HILIC | Neg | -0.6848 | mRNA | -0.6502 | Activity | TRUE |
| 15 | 3 | 1 | Phenyl sulfate | 172.9909 | 1.27 | Liver | HILIC | Neg | -0.6914 | mRNA | -0.6766 | Activity | TRUE |
| 16 | 5 | 1 | Phenyl sulfate | 172.9906 | 1.57 | Plasma | RPLC | Neg | -0.6902 | mRNA | 0 | 0 | TRUE |
| 17 | 5 | 1 | Phenyl sulfate | 172.9906 | 1.59 | Liver | RPLC | Neg | **-0.726** | mRNA | -0.6844 | Activity | TRUE |
| 18 | 1 | 2 | 2-Octenoylcarnitine | 286.2011 | 1.82 | Liver | HILIC | Pos | **-0.8085** | **mRNA**/Protein/ Activity | 0 | 0 | TRUE |
| 19 | 0 | 2 | 2-Octenoylcarnitine | 286.2014 | 1.88 | Liver | RPLC | Pos | **-0.8225** | **mRNA**/Protein/ Activity | 0 | 0 | TRUE |

| | | | | | | | | | | | | | |
|---|---|---|---|---|---|---|---|---|---|---|---|---|---|
| 20 | 2 | 2 | 2-Phenylethanol glucuronide | 297.0973 | 1.91 | Liver | RPLC | Neg | **-0.7038** | mRNA/**Activity** | 0 | 0 | TRUE |
| 21 | 1 | 1 | Creatine | 128.0817 | 3.48 | Liver | HILIC | Pos | **-0.7243** | mRNA/**Activity** | 0 | 0 | FALSE |
| 22 | 1 | 2 | Daidzein | 255.0650 | 2.12 | Liver | RPLC | Pos | -0.6562 | Protein | 0 | 0 | FALSE |
| 23 | 0 | 2 | Daidzein | 255.0651 | 2.80 | Plasma | HILIC | Pos | -0.68 | Protein | 0 | 0 | FALSE |
| 24 | 0 | 2 | Daidzein | 255.0653 | 0.96 | Liver | HILIC | Pos | -0.6553 | Protein | 0 | 0 | TRUE |
| 25 | 1 | 2 | Daidzein | 255.0654 | 1.04 | Plasma | HILIC | Pos | -0.6503 | mRNA | 0 | 0 | FALSE |
| 26 | 0 | 2 | Daidzein-4/7-O-glucuronide | 431.0971 | 2.80 | Plasma | HILIC | Pos | **-0.7188** | mRNA/**Protein** | 0 | 0 | FALSE |
| 27 | 0 | 2 | Indole (M+H) | 118.0651 | 1.25 | Plasma | HILIC | Pos | **-0.7407** | **mRNA**/Protein/ Activity | -0.6618 | Activity | TRUE |
| 28 | 0 | 2 | Indoleacrylic Acid (M+H) | 188.0706 | 1.36 | Plasma | RPLC | Pos | 0 | 0 | **0.7221** | mRNA/**Activity** | TRUE |
| 29 | 3 | 2 | *LysoPE [18:1(11Z)/0:0 or 0.0/18:1(9Z) or 18.1 (9Z)/0:0 or 0:0/18.1(9Z)]* | 478.2927 | 4.06 | Liver | RPLC | Neg | -0.6749 | Activity | 0 | 0 | FALSE |
| 30 | 3 | 2 | O-methoxycatechol-O-sulphate (M-H) | 203.0014 | 1.61 | Liver | RPLC | Neg | **-0.7238** | mRNA | 0 | 0 | FALSE |
| 31 | 2 | 2 | p-Toluenesulfonic acid (M+FA-H) or Tyrosol-4-sulfate (M-H) | 217.0172 | 1.98 | Liver | HILIC | Neg | -0.6616 | Activity | 0 | 0 | TRUE |
| 32 | 4 | 2 | Pyrocatechol sulfate, catechol sulfate | 188.9856 | 1.54 | Liver | RPLC | Neg | -0.6734 | mRNA | -0.6615 | Activity | TRUE |
| 33 | 3 | 2 | Pyrocatechol sulfate, catechol sulfate (M-H) | 188.9857 | 0.85 | Liver | HILIC | Neg | -0.6857 | mRNA | 0 | 0 | TRUE |
| 34 | 0 | 2 | Tyramine-O-sulfate (M+H) | 218.0481 | 2.68 | Plasma | HILIC | Pos | **-0.7758** | mRNA | **-0.7745** | mRNA/**Activity** | FALSE |
| 35 | 1 | 2 | Tyrosol-4-sulfate (M-H2O-H) | 199.0064 | 0.86 | Liver | HILIC | Neg | **-0.7261** | **mRNA**/Activity | 0 | 0 | TRUE |
| 36 | 1 | 2 | Tyrosol-4-sulfate (M-H2O-H) | 199.0064 | 2.01 | Plasma | RPLC | Neg | **-0.7414** | **mRNA**/Activity | 0 | 0 | TRUE |
| 37 | 0 | 2 | Tyrosol-4-sulfate (M-H2O-H) | 199.0065 | 2.04 | Liver | RPLC | Neg | **-0.7029** | mRNA | -0.6734 | Activity | TRUE |
| 38 | 0 | 2 | Tyrosol-4-sulfate (M-H2O-H) | 199.0065 | 1.03 | Liver | HILIC | Neg | **-0.7538** | **mRNA**/Activity | 0 | 0 | TRUE |
| 39 | 0 | 3 | *PC, PE or DG (M+H)* | 780.5538 | 2.56 | Plasma | HILIC | Pos | -0.674 | Activity | 0 | 0 | FALSE |
| 40 | 2 | 3 | (23S)-23,25-dihdroxy-24-oxovitamine D3 23-(beta-glucuronide) (M+K-2H) | 643.2878 | 4.39 | Liver | RPLC | Neg | 0 | 0 | 0.6977 | mRNA/**Protein** | TRUE |
| 41 | 0 | 3 | (23S)-23,25-dihdroxy-24-oxovitamine D3 23-(beta-glucuronide) (M+K-2H) | 643.2892 | 3.51 | Liver | HILIC | Neg | 0 | 0 | 0.6637 | Protein | FALSE |
| 42 | 1 | 3 | 2'-Deoxyinosine triphosphate (M+Na-2H) | 512.9600 | 0.80 | Liver | HILIC | Neg | 0 | 0 | **0.794** | mRNA/Protein/ **Activity** | FALSE |
| 43 | 1 | 3 | 3-Methylene-indolenine (M+H) | 130.0650 | 1.96 | Plasma | HILIC | Pos | -0.6943 | mRNA | 0 | 0 | FALSE |
| 44 | 0 | 3 | 3-Sulfodeoxycholic acid (M+H+K) | 249.1021 | 2.18 | Plasma | HILIC | Pos | -0.6533 | mRNA | 0 | 0 | TRUE |
| 45 | 1 | 3 | 4-Guanidinobutanoic acid (M+H) | 146.0925 | 3.47 | Liver | HILIC | Pos | **-0.7097** | mRNA/**Activity** | 0 | 0 | TRUE |

| | | | | | | | | | | | | | |
|---|---|---|---|---|---|---|---|---|---|---|---|---|---|
| 46 | 3 | 3 | 4-Guanidinobutanoic acid (M+H) | 146.0919 | 0.76 | Plasma | RPLC | Pos | -0.6606 | Activity | 0 | 0 | FALSE |
| 47 | 2 | 3 | 7'-Carboxy-alpha-chromanol (M+H) | 349.2365 | 1.92 | Liver | RPLC | Pos | -0.6684 | Protein | **-0.7467** | **mRNA**/Protein/ Activity | FALSE |
| 48 | 1 | 3 | 9-Decenoylcarnitine (M+H) | 314.2322 | 1.71 | Liver | HILIC | Pos | -0.6582 | mRNA | 0 | 0 | FALSE |
| 49 | 3 | 3 | Adenosine phosphosulfate (M+Na-2H) | 447.9960 | 2.07 | Plasma | HILIC | Neg | -0.6885 | mRNA | 0 | 0 | FALSE |
| 50 | 2 | 3 | *Androgen Derivitive (M+H+Na)* | 158.1180 | 0.71 | Liver | RPLC | Pos | **0.7429** | Activity | 0 | 0 | TRUE |
| 51 | 0 | 3 | *Cardiolipin* (M-3H) | 513.3115 | 0.88 | Liver | HILIC | Neg | 0.6704 | mRNA | **0.7833** | **mRNA**/Protein/ Activity | FALSE |
| 52 | 0 | 3 | Coumestrin or Daidzein 4/7-O-glucuronide (M-H) | 429.0827 | 2.79 | Liver | HILIC | Neg | **-0.7323** | Protein | 0 | 0 | FALSE |
| 53 | 1 | 3 | Cytidine triphosphate (M-H2O-H) | 463.9667 | 0.81 | Liver | HILIC | Neg | 0 | 0 | **0.7506** | mRNA/**Activity** | FALSE |
| 54 | 2 | 3 | Deoxycorticosterone (M+H) | 331.2274 | 0.85 | Liver | HILIC | Pos | 0 | 0 | -0.6813 | mRNA/**Activity** | FALSE |
| 55 | 3 | 3 | Dihydrowyerol (2M-H) | 523.2354 | 2.97 | Liver | RPLC | Neg | 0 | 0 | **-0.7006** | **mRNA**/Protein | FALSE |
| 56 | 0 | 3 | Dihyrdocortisol (M-H2O-H) | 345.2067 | 3.04 | Plasma | RPLC | Neg | **-0.7343** | mRNA/**Protein** | **-0.7892** | mRNA/Protein/ **Activity** | FALSE |
| 57 | 1 | 3 | Equol (M+H) | 243.1013 | 1.85 | Liver | RPLC | Pos | **-0.7218** | **mRNA**/Protein | **-0.7529** | mRNA/**Activity** | FALSE |
| 58 | 2 | 3 | *Estrone sulfate (M+2H) or Guanidinosuccinic acid (M+H)* | 176.0669 | 4.14 | Liver | HILIC | Pos | -0.6823 | Activity | 0 | 0 | FALSE |
| 59 | 1 | 3 | Glycerylphosphorylethanolamine (M+H) | 216.0634 | 4.78 | Liver | HILIC | Pos | 0 | 0 | 0.6938 | mRNA/**Activity** | TRUE |
| 60 | 2 | 3 | Glycerylphosphorylethanolamine (M-H) | 214.0481 | 4.82 | Liver | HILIC | Neg | 0 | 0 | 0.6562 | Activity | TRUE |
| 61 | 4 | 3 | Homocitrulline | 188.1033 | 4.58 | Liver | HILIC | Neg | -0.6989 | **mRNA**/Activity | 0 | 0 | FALSE |
| 62 | 5 | 3 | Homocitrulline (M-H) | 188.1031 | 0.59 | Liver | RPLC | Neg | -0.6567 | mRNA | 0 | 0 | FALSE |
| 63 | 3 | 3 | L-Acetylcarnitine (M+H) | 204.1237 | 0.71 | Liver | RPLC | Pos | -0.6587 | Protein | 0 | 0 | FALSE |
| 64 | 2 | 3 | L-gamma-glutamyl-L-leucine or L-gamma-glutamyl-L-isoleucine (M-H) | 259.1294 | 2.50 | Liver | HILIC | Neg | -0.6569 | mRNA | 0 | 0 | FALSE |
| 65 | 0 | 3 | LysoPC (18:3) (M+FA-H) | 562.3149 | 3.33 | Plasma | RPLC | Neg | **-0.7105** | Activity | 0 | 0 | TRUE |
| 66 | 0 | 3 | LysoPC (18:3) (M+FA-H) | 562.3152 | 3.01 | Plasma | HILIC | Neg | -0.679 | Activity | 0 | 0 | FALSE |
| 67 | 0 | 3 | *LysoPE (0:0/20:0 or 20:0/0:0) or LysoPC (17:0 or 16:0)* | 554.3461 | 4.58 | Plasma | RPLC | Neg | -0.6534 | Activity | 0 | 0 | TRUE |
| 68 | 0 | 3 | Nonacosanoic acid (M+2Na) | 242.2111 | 2.74 | Liver | HILIC | Pos | **-0.8032** | **mRNA**/Activity | -0.6673 | Activity | TRUE |
| 69 | 0 | 3 | Palmitic acid (M+2Na-H) | 301.2113 | 3.55 | Liver | HILIC | Pos | 0.657 | Activity | 0 | 0 | FALSE |
| 70 | 0 | 3 | *PC or PE* | 800.5448 | 2.61 | Liver | HILIC | Neg | -0.6577 | Activity | 0 | 0 | TRUE |
| 71 | 2 | 3 | Perfluorooctanesulfonic acid (M-H) | 498.9292 | 4.88 | Liver | RPLC | Neg | 0 | 0 | **0.8092** | mRNA/**Activity** | TRUE |
| 72 | 1 | 3 | Perfluorooctanesulfonic acid (M-H) | 498.9297 | 0.53 | Liver | HILIC | Neg | 0 | 0 | **0.7866** | mRNA/**Activity** | TRUE |

| | | | | | | | | | | | | | |
|---|---|---|---|---|---|---|---|---|---|---|---|---|---|
| 73 | 0 | 3 | Phloroacetophenone 6'-[xylosyl-(1->6)-glucoside] (M-H2O-H) | 471.1503 | 0.52 | Liver | RPLC | Neg | 0 | 0 | **0.7089** | mRNA/Protein/ **Activity** | FALSE |
| 74 | 0 | 3 | Propyl-Tryptophan or Tryptophyl-Proline (M+NH4) | 319.1766 | 1.47 | Liver | RPLC | Pos | 0 | 0 | **0.8628** | mRNA/Protein/ **Activity** | FALSE |
| 75 | 2 | 3 | S-Adenosylhomocysteine (M+IsoProp+H) | 445.1876 | 1.71 | Liver | RPLC | Pos | **-0.7138** | **mRNA**/Protein | **-0.8839** | **mRNA**/Protein/ Activity | FALSE |
| 76 | 1 | 3 | Taurine (M+2Na+H) | 169.9857 | 3.94 | Plasma | HILIC | Pos | -0.6583 | Activity | 0 | 0 | FALSE |
| 77 | 1 | 3 | *Testosterone, Epitestosterone or Dehydroepiandrosterone (M+FA-H) sulfates* | 413.1636 | 2.18 | Liver | HILIC | Neg | -0.6896 | mRNA/**Protein** | **-0.7731** | **mRNA**/Protein/ Activity | TRUE |
| 78 | N/A | N/A | Unknown | 93.0341 | 1.27 | Liver | HILIC | Neg | -0.6844 | mRNA | -0.67 | Activity | TRUE |
| 79 | N/A | N/A | Unknown | 114.0301 | 0.56 | Liver | RPLC | Neg | **-0.7222** | mRNA | **-0.7811** | mRNA/**Activity** | FALSE |
| 80 | N/A | N/A | Unknown | 121.0282 | 2.23 | Plasma | HILIC | Pos | -0.671 | **mRNA**/Activity | -0.6651 | Activity | FALSE |
| 81 | N/A | N/A | Unknown | 122.0366 | 0.83 | Plasma | HILIC | Neg | -0.6599 | mRNA | 0 | 0 | FALSE |
| 82 | N/A | N/A | Unknown | 132.0442 | 1.07 | Plasma | HILIC | Pos | -0.6962 | **mRNA**/Activity | 0 | 0 | TRUE |
| 83 | N/A | N/A | Unknown | 138.0547 | 0.58 | Plasma | RPLC | Pos | **-0.7194** | mRNA | 0 | 0 | TRUE |
| 84 | N/A | N/A | Unknown | 138.0550 | 3.70 | Liver | HILIC | Pos | -0.6984 | mRNA | 0 | 0 | TRUE |
| 85 | N/A | N/A | Unknown | 151.0612 | 2.39 | Plasma | HILIC | Pos | **-0.7681** | mRNA | **-0.8363** | mRNA/**Activity** | FALSE |
| 86 | N/A | N/A | Unknown | 158.1174 | 0.79 | Plasma | RPLC | Pos | 0.6553 | Activity | 0 | 0 | TRUE |
| 87 | N/A | N/A | Unknown | 166.0361 | 4.60 | Plasma | HILIC | Neg | 0 | 0 | -0.676 | Activity | FALSE |
| 88 | N/A | N/A | Unknown | 167.0926 | 3.29 | Plasma | HILIC | Pos | 0 | 0 | -0.6574 | Activity | FALSE |
| 89 | N/A | N/A | Unknown | 170.0924 | 5.30 | Plasma | HILIC | Pos | -0.6745 | mRNA | 0 | 0 | TRUE |
| 90 | N/A | N/A | Unknown | 170.0924 | 5.37 | Liver | HILIC | Pos | -0.653 | mRNA | 0 | 0 | FALSE |
| 91 | N/A | N/A | Unknown | 173.9936 | 1.11 | Liver | HILIC | Neg | **-0.7255** | mRNA | -0.6642 | Activity | TRUE |
| 92 | N/A | N/A | Unknown | 181.0497 | 1.41 | Liver | RPLC | Neg | **-0.705** | Protein | 0 | 0 | FALSE |
| 93 | N/A | N/A | Unknown | 183.0403 | 3.52 | Liver | HILIC | Neg | **-0.7987** | **mRNA**/Protein/ Activity | -0.6912 | mRNA/**Activity** | FALSE |
| 94 | N/A | N/A | Unknown | 201.1901 | 1.21 | Plasma | HILIC | Neg | -0.6626 | Activity | **0** | 0 | TRUE |
| 95 | N/A | N/A | Unknown | 204.0662 | 1.89 | Plasma | RPLC | Neg | -0.6858 | Activity | 0 | 0 | FALSE |
| 96 | N/A | N/A | Unknown | 205.1261 | 0.79 | Plasma | RPLC | Pos | 0 | 0 | **0.7301** | **mRNA**/Protein/ Activity | FALSE |
| 97 | N/A | N/A | Unknown | 210.0755 | 2.19 | Plasma | HILIC | Pos | -0.6554 | mRNA | **0** | 0 | FALSE |
| 98 | N/A | N/A | Unknown | 218.1538 | 0.75 | Liver | HILIC | Pos | 0 | 0 | 0.6908 | Activity | FALSE |
| 99 | N/A | N/A | Unknown | 224.8548 | 1.85 | Plasma | HILIC | Neg | 0 | 0 | **0.726** | **mRNA**/Activity | FALSE |
| 100 | N/A | N/A | Unknown | 227.9966 | 1.20 | Plasma | RPLC | Neg | -0.6756 | Activity | 0 | 0 | FALSE |
| 101 | N/A | N/A | Unknown | 230.0634 | 0.74 | Plasma | HILIC | Pos | -0.6873 | mRNA | 0 | 0 | TRUE |
| 102 | N/A | N/A | Unknown | 231.0665 | 0.75 | Plasma | HILIC | Pos | **-0.7236** | mRNA | -0.6923 | Activity | FALSE |

| | | | | | | | | | | | | | |
|---|---|---|---|---|---|---|---|---|---|---|---|---|---|
| 103 | N/A | N/A | Unknown | 231.0761 | 1.00 | Plasma | HILIC | Pos | **-0.7548** | mRNA | **-0.7845** | mRNA/**Activity** | FALSE |
| 104 | N/A | N/A | Unknown | 231.0767 | 1.79 | Liver | RPLC | Neg | **-0.7257** | **mRNA**/Activity | 0 | 0 | FALSE |
| 105 | N/A | N/A | Unknown | 231.0769 | 1.79 | Plasma | RPLC | Neg | -0.6786 | mRNA | 0 | 0 | TRUE |
| 106 | N/A | N/A | Unknown | 231.0770 | 1.80 | Liver | HILIC | Neg | **-0.7422** | **mRNA**/Protein | 0 | 0 | FALSE |
| 107 | N/A | N/A | Unknown | 233.0269 | 0.64 | Plasma | HILIC | Neg | **-0.7293** | mRNA | 0 | 0 | FALSE |
| 108 | N/A | N/A | Unknown | 240.9999 | 2.07 | Plasma | HILIC | Neg | -0.6938 | mRNA | 0 | 0 | FALSE |
| 109 | N/A | N/A | Unknown | 241.7574 | 0.55 | Plasma | RPLC | Neg | 0 | 0 | **0.7186** | mRNA/**Activity** | FALSE |
| 110 | N/A | N/A | Unknown | 245.1605 | 3.32 | Plasma | HILIC | Pos | -0.6985 | mRNA | -0.6949 | mRNA/**Activity** | FALSE |
| 111 | N/A | N/A | Unknown | 246.0740 | 2.08 | Liver | RPLC | Neg | **-0.706** | mRNA/**Protein** | **-0.7278** | mRNA/Protein/ **Activity** | FALSE |
| 112 | N/A | N/A | Unknown | 253.0492 | 2.41 | Liver | HILIC | Neg | -0.6641 | Protein | 0 | 0 | FALSE |
| 113 | N/A | N/A | Unknown | 259.9984 | 1.09 | Plasma | RPLC | Neg | 0.6681 | Activity | 0 | 0 | FALSE |
| 114 | N/A | N/A | Unknown | 265.0970 | 0.92 | Plasma | HILIC | Pos | **-0.715** | **mRNA**/Protein/ Activity | 0 | 0 | TRUE |
| 115 | N/A | N/A | Unknown | 275.0591 | 2.01 | Plasma | HILIC | Neg | **-0.7034** | **mRNA**/Protein | 0 | 0 | FALSE |
| 116 | N/A | N/A | Unknown | 276.0626 | 1.95 | Liver | HILIC | Neg | -0.6991 | mRNA/**Protein** | 0 | 0 | FALSE |
| 117 | N/A | N/A | Unknown | 277.1432 | 0.85 | Plasma | HILIC | Pos | -0.6755 | mRNA | -0.6888 | mRNA | FALSE |
| 118 | N/A | N/A | Unknown | 282.9917 | 0.63 | Liver | HILIC | Neg | **-0.7728** | **mRNA**/Protein/ Activity | **-0.7072** | mRNA/**Activity** | FALSE |
| 119 | N/A | N/A | Unknown | 288.9650 | 2.16 | Plasma | HILIC | Neg | -0.6687 | mRNA | 0 | 0 | FALSE |
| 120 | N/A | N/A | Unknown | 289.1113 | 2.18 | Liver | HILIC | Neg | **-0.7895** | **mRNA**/Protein | **-0.7199** | **mRNA**/Activity | FALSE |
| 121 | N/A | N/A | Unknown | 298.1005 | 1.91 | Liver | RPLC | Neg | -0.6875 | Activity | 0 | 0 | FALSE |
| 122 | N/A | N/A | Unknown | 298.9770 | 0.72 | Liver | HILIC | Neg | 0 | 0 | 0.6634 | mRNA | FALSE |
| 123 | N/A | N/A | Unknown | 305.1751 | 0.95 | Liver | HILIC | Neg | **0.7851** | **mRNA**/Protein | **0.7375** | **mRNA**/Activity | FALSE |
| 124 | N/A | N/A | Unknown | 310.1405 | 2.39 | Liver | HILIC | Neg | 0 | 0 | -0.6598 | **mRNA**/Protein | FALSE |
| 125 | N/A | N/A | Unknown | 315.0007 | 0.84 | Liver | HILIC | Neg | 0 | 0 | **-0.7257** | Activity | FALSE |
| 126 | N/A | N/A | Unknown | 316.1755 | 1.25 | Liver | RPLC | Pos | **-0.7758** | **mRNA**/Protein/ Activity | 0 | 0 | TRUE |
| 127 | N/A | N/A | Unknown | 317.1246 | 1.40 | Liver | RPLC | Neg | 0.6754 | Protein | **0.8004** | **mRNA**/Protein/ Activity | FALSE |
| 128 | N/A | N/A | Unknown | 319.1760 | 3.22 | Liver | HILIC | Pos | 0 | 0 | **0.8146** | mRNA/Protein/ **Activity** | FALSE |
| 129 | N/A | N/A | Unknown | 323.0583 | 1.88 | Liver | RPLC | Neg | 0 | 0 | -0.6504 | Activity | FALSE |
| 130 | N/A | N/A | Unknown | 323.9994 | 1.91 | Liver | RPLC | Neg | **-0.742** | mRNA | **-0.7192** | mRNA/**Activity** | FALSE |
| 131 | N/A | N/A | Unknown | 329.0689 | 2.01 | Liver | RPLC | Neg | **-0.7388** | **mRNA**/Protein | **-0.7346** | mRNA/**Activity** | FALSE |
| 132 | N/A | N/A | Unknown | 329.0698 | 2.06 | Liver | HILIC | Neg | **-0.7452** | **mRNA**/Protein | -0.6795 | **mRNA**/Activity | TRUE |
| 133 | N/A | N/A | Unknown | 330.2252 | 1.70 | Liver | RPLC | Pos | -0.6748 | Activity | 0 | 0 | FALSE |
| 134 | N/A | N/A | Unknown | 333.0064 | 1.91 | Liver | RPLC | Neg | -0.6588 | mRNA | 0 | 0 | TRUE |

| | | | | | | | | | | | | | |
|---|---|---|---|---|---|---|---|---|---|---|---|---|---|
| 135 | N/A | N/A | Unknown | 334.0096 | 1.91 | Liver | RPLC | Neg | -0.662 | mRNA | 0 | 0 | FALSE |
| 136 | N/A | N/A | Unknown | 334.0104 | 2.07 | Liver | HILIC | Neg | -0.657 | mRNA/**Protein** | 0 | 0 | FALSE |
| 137 | N/A | N/A | Unknown | 339.0540 | 1.95 | Liver | HILIC | Neg | -0.6732 | Activity | 0 | 0 | TRUE |
| 138 | N/A | N/A | Unknown | 345.1551 | 2.21 | Liver | HILIC | Neg | **-0.7491** | Activity | 0 | 0 | FALSE |
| 139 | N/A | N/A | Unknown | 345.2424 | 4.63 | Liver | RPLC | Neg | **0.7401** | **mRNA**/Protein/ Activity | 0 | 0 | FALSE |
| 140 | N/A | N/A | Unknown | 346.2095 | 3.06 | Liver | RPLC | Neg | **-0.711** | **mRNA**/Protein | **-0.7988** | **mRNA**/Protein/ Activity | FALSE |
| 141 | N/A | N/A | Unknown | 346.2736 | 0.69 | Liver | HILIC | Pos | 0 | 0 | 0.67 | mRNA | FALSE |
| 142 | N/A | N/A | Unknown | 347.2210 | 3.07 | Liver | RPLC | Pos | **-0.7363** | **mRNA**/Protein | **-0.8455** | **mRNA**/Protein/ Activity | TRUE |
| 143 | N/A | N/A | Unknown | 348.2372 | 1.36 | Liver | RPLC | Pos | 0 | 0 | **0.7313** | mRNA/Protein/ **Activity** | FALSE |
| 144 | N/A | N/A | Unknown | 351.0558 | 0.56 | Liver | RPLC | Neg | **-0.7333** | **mRNA**/Protein | **-0.7218** | mRNA/**Activity** | FALSE |
| 145 | N/A | N/A | Unknown | 363.2171 | 2.46 | Plasma | RPLC | Neg | -0.6630 | Protein | **-0.7957** | **mRNA**/Protein/ Activity | FALSE |
| 146 | N/A | N/A | Unknown | 372.1896 | 2.40 | Liver | RPLC | Neg | -0.6892 | mRNA | 0 | 0 | FALSE |
| 147 | N/A | N/A | Unknown | 372.2374 | 3.08 | Plasma | HILIC | Pos | **-0.7488** | **mRNA**/Activity | 0 | 0 | FALSE |
| 148 | N/A | N/A | Unknown | 377.0697 | 2.40 | Liver | HILIC | Neg | 0 | 0 | -0.6544 | mRNA | FALSE |
| 149 | N/A | N/A | Unknown | 377.1961 | 0.62 | Liver | HILIC | Neg | **-0.7503** | **mRNA**/Protein | 0 | 0 | FALSE |
| 150 | N/A | N/A | Unknown | 383.1123 | 1.85 | Liver | RPLC | Pos | 0 | 0 | **-0.7391** | mRNA/**Activity** | FALSE |
| 151 | N/A | N/A | Unknown | 385.1679 | 2.23 | Liver | RPLC | Neg | -0.6681 | mRNA/**Protein** | **-0.7956** | **mRNA**/Protein/ Activity | FALSE |
| 152 | N/A | N/A | Unknown | 397.1677 | 2.90 | Liver | RPLC | Neg | 0 | 0 | **0.7725** | mRNA/Protein/ **Activity** | FALSE |
| 153 | N/A | N/A | Unknown | 397.1680 | 2.50 | Liver | RPLC | Neg | 0 | 0 | **-0.7494** | **mRNA**/Protein/ Activity | FALSE |
| 154 | N/A | N/A | Unknown | 398.1711 | 2.90 | Liver | RPLC | Neg | 0 | 0 | **0.7103** | mRNA/Protein/ **Activity** | FALSE |
| 155 | N/A | N/A | Unknown | 399.1835 | 3.06 | Liver | RPLC | Neg | 0 | 0 | 0.6785 | Activity | FALSE |
| 156 | N/A | N/A | Unknown | 415.2480 | 0.58 | Plasma | HILIC | Neg | -0.6569 | Activity | 0 | 0 | FALSE |
| 157 | N/A | N/A | Unknown | 419.9772 | 0.84 | Liver | HILIC | Neg | 0 | 0 | 0.6928 | **mRNA**/Activity | FALSE |
| 158 | N/A | N/A | Unknown | 438.8927 | 0.86 | Liver | HILIC | Neg | 0 | 0 | 0.6722 | mRNA/**Activity** | FALSE |
| 160 | N/A | N/A | Unknown | 445.1891 | 1.91 | Liver | RPLC | Neg | -0.6805 | mRNA/**Protein** | **-0.8438** | **mRNA**/Protein/ Activity | TRUE |
| 161 | N/A | N/A | Unknown | 445.1899 | 2.42 | Liver | HILIC | Neg | 0 | 0 | **-0.7918** | **mRNA**/Protein/ Activity | TRUE |
| 163 | N/A | N/A | Unknown | 461.1840 | 1.70 | Liver | RPLC | Neg | **-0.7138** | **mRNA**/Protein | **-0.8925** | **mRNA**/Protein/ Activity | TRUE |
| 164 | N/A | N/A | Unknown | 461.1851 | 1.69 | Plasma | RPLC | Neg | **-0.7048** | mRNA/**Protein** | **-0.8933** | **mRNA**/Protein/ Activity | FALSE |
| 165 | N/A | N/A | Unknown | 462.9631 | 0.81 | Liver | HILIC | Neg | 0 | 0 | **0.8195** | mRNA/Protein/ **Activity** | FALSE |

| | | | | | | | | | | | | | |
|---|---|---|---|---|---|---|---|---|---|---|---|---|---|
| 166 | N/A | N/A | Unknown | 472.1533 | 0.51 | Liver | RPLC | Neg | 0 | 0 | **0.7155** | mRNA/Protein/ **Activity** | FALSE |
| 167 | N/A | N/A | Unknown | 473.2567 | 2.41 | Liver | RPLC | Neg | **-0.7012** | **mRNA**/Protein | 0 | 0 | TRUE |
| 168 | N/A | N/A | Unknown | 480.2779 | 3.33 | Liver | HILIC | Pos | -0.6608 | Activity | 0 | 0 | TRUE |
| 169 | N/A | N/A | Unknown | 481.1537 | 0.84 | Liver | HILIC | Neg | **0.7445** | **mRNA**/Protein | **0.7975** | mRNA/**Activity** | FALSE |
| 170 | N/A | N/A | Unknown | 481.2809 | 3.36 | Liver | HILIC | Pos | -0.6614 | Activity | 0 | 0 | TRUE |
| 171 | N/A | N/A | Unknown | 481.2810 | 3.33 | Liver | HILIC | Pos | -0.6609 | Activity | 0 | 0 | TRUE |
| 172 | N/A | N/A | Unknown | 482.1570 | 0.84 | Liver | HILIC | Neg | **0.7422** | **mRNA**/Protein | **0.7957** | mRNA/Protein/ **Activity** | FALSE |
| 173 | N/A | N/A | Unknown | 499.9329 | 0.53 | Liver | HILIC | Neg | 0 | 0 | **0.8034** | mRNA/**Activity** | TRUE |
| 174 | N/A | N/A | Unknown | 500.9272 | 0.53 | Liver | HILIC | Neg | 0 | 0 | **0.7574** | mRNA/**Activity** | FALSE |
| 175 | N/A | N/A | Unknown | 501.2154 | 2.01 | Liver | RPLC | Neg | **0.7268** | **mRNA**/Protein | **0.8195** | **mRNA**/Protein/ Activity | TRUE |
| 176 | N/A | N/A | Unknown | 501.9298 | 0.54 | Liver | HILIC | Neg | 0 | 0 | 0.6618 | Activity | FALSE |
| 177 | N/A | N/A | Unknown | 503.2181 | 2.37 | Liver | HILIC | Neg | 0 | 0 | 0.6648 | mRNA | FALSE |
| 178 | N/A | N/A | Unknown | 505.2244 | 4.13 | Liver | RPLC | Neg | 0 | 0 | 0.6503 | mRNA | FALSE |
| 179 | N/A | N/A | Unknown | 507.2726 | 2.79 | Liver | HILIC | Neg | 0 | 0 | -0.6951 | Activity | TRUE |
| 180 | N/A | N/A | Unknown | 521.4929 | 0.67 | Plasma | HILIC | Neg | -0.6603 | mRNA | 0 | 0 | FALSE |
| 181 | N/A | N/A | Unknown | 523.9726 | 2.13 | Plasma | HILIC | Neg | -0.6503 | Protein | 0 | 0 | FALSE |
| 182 | N/A | N/A | Unknown | 528.1117 | 2.05 | Plasma | HILIC | Neg | -0.6803 | mRNA | 0 | 0 | FALSE |
| 183 | N/A | N/A | Unknown | 555.0567 | 1.14 | Plasma | HILIC | Neg | **0.7185** | mRNA | 0 | 0 | FALSE |
| 184 | N/A | N/A | Unknown | 561.3085 | 2.30 | Liver | RPLC | Neg | 0 | 0 | 0.662 | Activity | FALSE |
| 185 | N/A | N/A | Unknown | 593.2730 | 3.57 | Liver | HILIC | Neg | -0.6822 | Activity | 0 | 0 | FALSE |
| 186 | N/A | N/A | Unknown | 595.2879 | 4.35 | Liver | RPLC | Neg | -0.6675 | Activity | 0 | 0 | TRUE |
| 187 | N/A | N/A | Unknown | 595.2891 | 3.56 | Liver | HILIC | Neg | -0.6799 | Activity | 0 | 0 | TRUE |
| 189 | N/A | N/A | Unknown | 599.9271 | 0.53 | Liver | HILIC | Neg | 0 | 0 | 0.6541 | mRNA/**Activity** | FALSE |
| 190 | N/A | N/A | Unknown | 603.5334 | 3.15 | Liver | HILIC | Pos | -0.6776 | Activity | 0 | 0 | TRUE |
| 191 | N/A | N/A | Unknown | 609.3437 | 0.63 | Liver | HILIC | Neg | **-0.8106** | mRNA | **-0.7937** | mRNA/**Activity** | FALSE |
| 192 | N/A | N/A | Unknown | 610.3468 | 0.63 | Liver | HILIC | Neg | **-0.7263** | **mRNA**/Protein | -0.6784 | **mRNA**/Activity | FALSE |
| 193 | N/A | N/A | Unknown | 645.3033 | 4.75 | Liver | RPLC | Neg | 0 | 0 | **0.7562** | **mRNA**/Protein/ Activity | FALSE |
| 194 | N/A | N/A | Unknown | 645.3035 | 3.51 | Liver | HILIC | Neg | 0 | 0 | **0.7334** | mRNA/**Protein** | FALSE |
| 195 | N/A | N/A | Unknown | 647.3421 | 2.49 | Liver | RPLC | Neg | 0.6805 | mRNA | 0.6849 | **mRNA**/Activity | FALSE |
| 196 | N/A | N/A | Unknown | 651.3756 | 2.01 | Plasma | HILIC | Neg | **0.7103** | mRNA | 0.6664 | Activity | FALSE |
| 197 | N/A | N/A | Unknown | 714.2969 | 2.12 | Liver | RPLC | Neg | **-0.7594** | Activity | 0 | 0 | FALSE |
| 198 | N/A | N/A | Unknown | 785.3793 | 3.25 | Liver | HILIC | Pos | **-0.7231** | Activity | 0 | 0 | FALSE |
| 199 | N/A | N/A | Unknown | 864.5905 | 2.46 | Liver | HILIC | Pos | 0.6522 | mRNA | 0.6712 | Activity | TRUE |

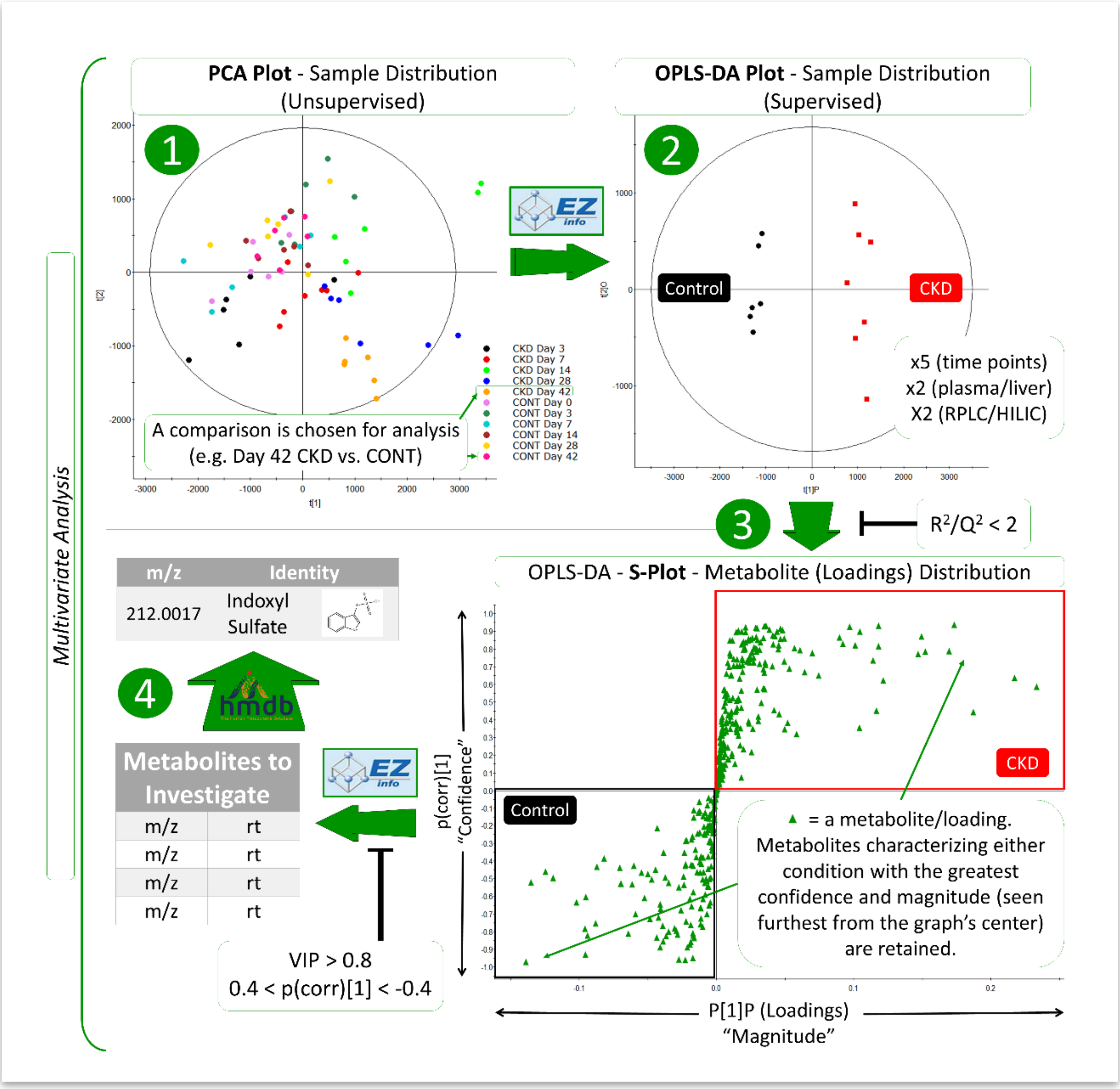


**Supplementary Figure 1.** Example of multivariate analysis workflow utilizing OPLS-DA and S- is made using EZInfo software (1), comparisons are chosen for OPLS-DA, an OPLS-DA pl distribution is made and $R^2$ and $Q^2$ thresholds are applied (2). Successful OPLS-DA plots are fur by observing metabolite contributions towards either condition by S-plot (3) where VIP a thresholds are applied. From the S-plot, a list of *m/z* ratios and retention times (4) in addition to pattern are searched within online databases such as HMDB in attempt to identify the metabolite.
